# Multiple transcription factors mediate acclimation of Chlamydomonas to light stress

**DOI:** 10.1101/2023.10.30.564712

**Authors:** Donat Wulf, Fabian Janosch Krüger, Levin Joe Klages, Prisca Viehöver, EonSeon Jin, Lutz Wobbe, Marion Eisenhut, Olaf Kruse, Olga Blifernez-Klassen, Andrea Bräutigam

## Abstract

Light as a substrate for photosynthesis may be a boon or a bane. To thrive, photosynthetic organisms must constantly respond to changing light and CO2 conditions by balancing energy harvest and consumption in a highly dynamic way. Two major safeguard measures of photoacclimation, that is photoprotection and carbon concentrating mechanism, underlie tight transcriptional control, leading to expression changes under high light and limited CO2 with different dynamics for both systems. Here, by using a consensus gene regulatory network inferred by employing a compendium of 1,869 RNA-seq datasets, we identified and validated *in vivo* eight candidate transcription factors (TFs) that contribute to photoacclimation in *Chlamydomonas reinhardtii*. Target gene analyses indicate that the TFs act individually in associated pathways but also influence each other in expression, and function as network parts with partial redundancy with respect to photoprotection. The analyses unveil that stress responses in *Chlamydomonas* are mediated by a complex, interconnected network of TFs rather than a hierarchical system where multiple regulators can influence each other and target gene expression and thereby mitigate the effects of loss.

## Introduction

Gene regulation in photosynthetic eukaryotes is complex. Each gene is controlled by its promoter and possible enhancers to which the general transcription machinery and sequence-specific transcription factors (TFs) bind (Liu et al., 1999). Enhancer elements can be in the first intron of a gene, the 5’ UTR (Stergachis et al., 2013; Gallegos and Rose, 2019; Rose, 2019), and several kb distant from the transcriptional start side (Schmitz et al., 2022). Responses to stressors, such as excess light, or metabolic pathways are controlled by one or several TFs (Song et al., 2016). In microalgae, such as *Chlamydomonas reinhardtii* the light is harvested by light-harvesting complexes (LHCs), which transfer absorbed energy to photosystem (PS) I and PSII. If light is harvested in excess, energy cannot be fully consumed by the Calvin Benson Bassham cycle (CBBc) and other energy-consuming cellular processes. As a result, electron acceptors get limited and excess electrons lead to the formation of toxic reactive oxygen species (ROS) (Niyogi, 1999). Many biotic and abiotic factors alter the balance between energy harvest and energy consumption making light protection a highly dynamic process. Light changes throughout the day in quantity and quality in a predictable manner (Thorne et al., 2009). Algae constantly face changes in CO2 supply (Provenzale et al., 2018) in the water column (Cory et al., 2014), which leads to changes in flux through the CBBc and therefore changes in electron acceptor regeneration. Algae also alter their position relative to light depending on water currents and their swimming, which again changes light quantity and quality (Erickson et al., 2015). The nightly division dilutes light protective proteins in half and requires replenishment upon or before the onset of light in the morning (Cross and Umen, 2015; Strenkert et al., 2019).

In *C. reinhardtii*, two main components of photoacclimation regulation are known: photoprotection (PP) itself and the carbon concentrating mechanism (CCM). PP is mediated via stress-related light-harvesting proteins, called LHCSRs that contribute to non-photochemical quenching and are required for survival in a dynamically changing environment (Peers et al., 2009). They act as quenchers on both photosystems (Girolomoni et al., 2019). PsbS also contributes to PP in *C. reinhardtii* (Redekop et al., 2020), although its molecular mode of action is currently unclear. An analysis of mutants, such as *npq4*, indicates that LHCSR3 is the major contributor to PP (Redekop et al., 2020). *C. reinhardtii* can also increase the regeneration of electron acceptors by concentrating CO2 through a CCM and thus boost the CBBc activity (Wang et al., 2015). High light and limited CO2 were demonstrated to induce changes in expression with different dynamics for both systems (Ruiz-Sola et al., 2023; Redekop et al., 2022; Tibiletti et al., 2016). A subset of the relevant inputs and the controlling TFs are known. Light inputs are transduced to PP gene expression via the E3 ubiquitin ligase CONSTITUTIVE PHOTOMORPHOGENIC 1 (COP1) as shown in the constitutively high-light acclimated *hit1-1* mutant (Schierenbeck et al., 2015). COP1 acts on PP genes via a homologue of the TF CONSTANS (CO) (Gabilly et al., 2019). The GARP-type TF RHYTHM OF CHLOROPLAST 75 (ROC75) also controls LHCSR expression (Kamrani et al., 2018). A UV light signal can be perceived through UV-B resistance 8 (UVR8)/COP1 (Tilbrook et al., 2016; Favory et al., 2009), and also controls LHCSRs (Allorent et al., 2016). The blue light signal induces expression of the PP gene LHCSR3 (Petroutsos et al., 2016) and of other photosynthetic transcripts (Im et al., 2006) via phototropin, but the mediating TFs are currently unknown. Red and blue light are perceived through the cryptochromes (CRYs) aCRY and pCRY (Reisdorph and Small, 2004; Beel et al., 2012). Phototropin and pCRY control the life cycle (Müller et al., 2017; Huang and Beck, 2003), and aCRY controls chlorophyll and carotenoid biosynthesis (Beel et al., 2012), indicating that light also affects pathways other than photoacclimation. Limiting CO2 supply rapidly activates the response factor LCRF and this TF triggers the accumulation of the nucleic acid binding protein NAB1, which in turn enhances the translational repression of PSII-associated light-harvesting proteins (LHCBMs) (Blifernez-Klassen et al., 2021). The CO2 inputs are transduced via the transcriptional regulator CIA5 (also named CCM1) to both CCM and PP gene expression (Fang et al., 2012) and via the myb-type TF LCR1 (for low-CO2 stress response) (Yoshioka et al., 2004; Arend et al., 2023). The TF qE-REGULATOR 7 (QER7) belonging to the SQUAMOSA-PROMOTER BINDING PROTEIN-LIKE gene family was identified via network analyses also controls both PP and CCM genes (Arend et al., 2023). Unlike many other TFs (Yoshioka et al., 2004; Arend et al., 2023; Tokutsu et al., 2019), CIA5 has been analyzed with global RNA-seq and shown to affect pathways other than PP and CCM (Fang et al., 2012). Other inputs, such as ABA, which increases carbon uptake in *C. reinhardtii* and alters swimming behaviour (Al-Hijab et al., 2019) as well as redox signals that change PP gene expression (Redekop et al., 2022), transduce their signal through unknown mechanisms. The unknown TFs for known input signals indicate substantial discovery potential for novel TFs. Furthermore, the limited analyses of single target genes prompt questions about the extent of the downstream response for photoacclimation TFs.

Several large to very large expression datasets pertaining to the induction of PP have been produced in *C. reinhardtii,* such as a diurnal dataset (Zones et al., 2015), CO2 concentration changes (Fang et al., 2012), and redox challenges (Ma et al., 2020). A curated set of expression data was successfully used to identify QER7, which controls the expression of PP genes (Arend et al., 2023). Co-expression analyses of the diurnal data have revealed that light stress gene expression peaks at the onset of light, at the sharp transition between dark and light together with re-flagellation while photosynthetic gene expression peaks in the middle of the day (Zones et al., 2015). These datasets and additional FAIR (findable, accessible, interoperable, and reusable) data in the sequence read archive theoretically enable large-scale co-expression analyses (Salomé and Merchant, 2021) and gene regulatory network construction (Ramírez-González et al., 2018; Halpape et al., 2023). Co-expression analysis assumes a linear relationship between the target genes and their controlling TF. In contrast, random-forest (RF) decision tree-based gene regulatory network construction is capable of representing linear, non-linear, and combinatorial effects (Breiman, 2001; Huynh-Thu et al., 2010).

Here we present the identification and characterization of a network of eight TFs and the ubiquitin ligase COP1 that contribute to photoacclimation in *C. reinhardtii* (Schierenbeck et al., 2015). We initially used a RNA-seq data compilation of 769 datasets to test if expression patterns of PP and CCM transcripts converge with known regulators. An RF-based approach then identified new candidate TFs, for eight of which knock-out mutants were established and characterized with regard to growth, NPQ performance, and transcriptome. DEG analyses of the mutants showed that photoacclimation likely includes not only photoacclimation genes but also upregulation of photorespiration and downregulation of light harvesting as well as changes in expression for ciliogenesis and DNA damage control. The TFs indeed act as network parts with partial redundancy in photoacclimation and they act individually in associated pathways. The data reveals that stress responses in *C. reinhardtii* are mediated through a complex, interconnected network of TFs that act directly and indirectly on their target genes and integrate various inputs into a common response.

## Material and Methods

Data analysis was performed with R 4.3.1 and python 3.9.17.

### GENIE3-based gene regulatory network construction and comparison to correlation-based network

The sequence read archive (Katz et al., 2022) provided 1,868 RNA-seq datasets (“*C. reinhardtii”* AND “RNA”). The dataset was manually curated for wild-type transcriptomic studies using mRNA (Supplemental Table 1, 769 datasets). The data were downloaded with fasterq-dump (v2.11.2) and mapped on the primary transcriptome of *C. reinhardtii* v.5.6 Kallisto (v0.46.1) (Bray et al., 2016). The resulting expression matrix is publicly available as data publication (https://gitlab.ub.uni-bielefeld.de/computationalbiology/cr_grn/-/blob/main/wild-type_RNA-seq_expression_matrix.tsv.gz) Expression matrix was z-scaled gene wise and clustered by the Euclidian distance with complete linkage in gene and experiment direction.

237 *C. reinhardii* TFs were obtained from TAP-scan (Petroll et al., 2021) (Supplemental Table 2). To generate candidate TF – target pathway associations, the random forest decision tree regression based algorithm GENIE3 (Huynh-Thu et al., 2010) was used to calculate the weighted connections of 237 TFs with 17,741 target genes using the expression matrix (https://gitlab.ub.uni-bielefeld.de/computationalbiology/cr_grn/-/blob/main/wild-type_RNA-seq_expression_matrix.tsv.gz) with 1,000 decision trees and the square root of regulators. For the correlation-based network the expression matrix was z-scaled and the Pearson correlation coefficient was calculated for each TF and all other genes For the correlation based network cutoffs between 0 and 1 in 75 steps and for the GENIE3 network for cutoffs from 0.6/1.2^x^ (x=[1-75]) were used as sliding cut-offs. Gene Ontology terms were transferred based on the best blastp (Altschul et al., 1990) hit from *A. thaliana*. *C. reinhardtii* genes were assigned to *A. thaliana* TAIR10 genes by a blastp search (e<1^-5^). Genes with a weight or correlation coefficient above the cut-off were defined as target genes and a GOterm enrichment was performed with topGO (v2.52.0) (Alexa and Rahnenführer) using the classic fisher method for the ontology biological process. To homogenize the data range for plotting the weight cut-off of the GENIE3 was scaled log10 and then rescaled between 0 and 1. To identify enrichments in particular pathways without well annotated gene ontology, the photosynthesis, CBBc, and LHC lists were derived from KEGG (Kanehisa et al., 2017). The CCM gene list was transferred from Mallikarjuna et al., 2020. The PP list was manually curated (Supplement Table 3). For each gene list, a fisher exact test was performed to test for a significant enrichment in predicted target genes of the GENIE3 network with a network cut-off of 0.005.

### Strains and Growth conditions

All microalgal strains were grown photoautotrophically in high salt medium HSM (Sueoka et al., 1967). Low light – high CO2 conditions (LL conditions) refer to 70 - 80 µmol photons s^-1^ m^-2^ and air enriched with CO2 (5 % CO2 [v/v]), while high light – low CO2 conditions refer to 350 - 400 µmol photons s^-1^ m^-2^ and regular atmospheric air (0.04% CO2 [v/v]) (HL conditions).

### Validation of insertion sites in CLiP mutants

Genomic DNA was isolated using the Quick-DNA Miniprep kit (Zymo Research) according to manufacturer instructions. For nanopore long read sequencing, the ligation sequencing kit with native barcodes (SQK-LSK114 with SQK-NBD114.24) was used according to the manufacturer’s protocol (Oxford Nanopore Technology). Long read data were produced on a R10.4.1 flowcell (Oxford Nanopore) which was run on a GridION X5 sequencing platform and base called using guppy (Wick et al., 2019) v7.0.9 in super-accuracy mode. Reads were mapped with minimap2 (Li, 2018) (2.26-r1175) against the CLiP insertion cassette and filtered for a mapping quality greater than 10 and an overlap with the cassette in the first 50 bp and the last 63 bp with samtools (Danecek et al., 2021) (v1.18) because these regions are not part of the nuclear encoded genome of *C. reinhardtii* (v5.6). These reads were than mapped on the genome of *C. reinhardtii* to confirm the insertion site and to check for additional insertion sites. For visualization of the insertions the identified reads were aligned with blastn with the insertion position. The insertion position left and right from the cassette was determined by identification of common start and end positions of the reads. The genomic sequence was determined by concatenating the genomic sequence and the cassette based on the identified insertion position. The reads were aligned with blastn to the combination of genomic sequence and the cassette to determine the coverage. Exons in the region were aligned with blastn to the genomic region including the insertion cassette. The coverage, the identified genomic context and the exons were visualised with gggenomes (v0.9.9.9) (Hackl et al.). Additionally, the whole sample was mapped with minimap2 to the genome of *C. reinhardtii* (v5.6) to check that no read without the CLiP insertion cassette spanned the insertion site.

### Growth and nonphotochemical quenching measurements

To assess the growth behavior of the mutants and wildtype, LL-preadapted cells were cultivated for 4 days in constant white light (Osram L 36 W/865, Osram, Germany) at 350 µmol photons s^-1^ m^-2^ and gassed with air enriched with CO2 (5 % CO2 [v/v], ML-conditions). The increase in cell density was monitored by cell counts using a haemocytometer (Neubauer Improved). Specific growth rates *μ* were calculated for the exponential phase using the following equation, where *N*2 represents the cell number per ml at *t*2 and *N*1 the respective number at *t*1: µ = ln(N2/N1)/(t2-t1). The students t-test determined significant differences (n=3).

For fluorescence analyses, LL-preadapted algae cultures were adjusted to a chlorophyll concentration of ∼7.5 µg Chl/mL as described previously (Berger et al., 2014) and incubated for 2h at HL-conditions and used for fluorescence analyses. Prior to chlorophyll fluorescence measurements, cells suspensions were acclimated to darkness for 2 h. NPQ was calculated as (Fm-Fm’)/Fm’ using FluorCam FC 800-C video imaging system (Photon system instruments, Drasov, Czech Republic) in a 48-well plate (flat bottom) in 3 biological and 5 technical replicates. Fm is the maximal Chl fluorescence emitted of dark-adapted cells after 2 min treatment of far-red light and Fm’ is the maximal fluorescence yield recorded during actinic light illumination of 1,100µmol m^-2^s^-1^ (Maxwell and Johnson, 2000). Significant differences were tested with a students t-test.

### RNA isolation, sequencing, and differential gene expression analysis

LL-preadapted microalgal cells were inoculated at an OD750 of 0.1 in three biological replicates and grown to an OD750 of 0.5 in LL conditions. They were subsequently moved to HL conditions and sampled by centrifugation for 5 min at 3,000 x *g* after 30 min. Total RNA was isolated using the Quick-RNA Plant Miniprep kit (Zymo Research) according to manufacturer instructions. RNA quality and quantity were assessed using the Bioanalyzer RNA 6000 Nano Kit (Agilent) following the instructions provided by the manufacturer. Samples scoring an RNA Integrity Number greater than 7.0 were used for further analysis. Library preparation and sequencing were carried out according to the instructions with the TruSeq RNA Library Prep Kit v2 (Illumina) and NextSeq 500/550 High Output Kit (Illumina). Reads were mapped as described and differential gene expression analysis was performed with edgeR (Robinson et al., 2010) (v3.42.4.). Genes with an FDR<0.01 were considered significantly differentially abundant. GO term enrichment was performed with topGO (Alexa and Rahnenführer) (v2.52.0) on significantly up and downregulated genes. Kegg Ontology (KO) numbers were obtained from the functional annotation v5.6 and assigned their pathways based on KEGG (Kanehisa et al., 2017) brite database (https://www.genome.jp/kegg-bin/get_htext accessed 25^th^ August 2023). Pathways were manually curated for pathways present in *C. reinhardii* (Supplement table 4). Significant enrichment was calculated with a fisheŕs exact test.

For Nitrogen deficiency DEG analysis was performed by comparing the control samples (SRX113670, SRX113655) against the 8 h samples (SRX113673, SRX113658) (Boyle et al., 2012). For iron limitation conditions, iron limited (SRR402024, SRR402025, SRR402026) samples were compared against iron deficient samples (SRR402021, SRR402022, SRR402023) (Urzica et al., 2012b). For the oxidative stress set, control samples (SRX113857) were compared to 1 h H2O2 treatment samples(SRX113859) (Urzica et al., 2012a). The low CO2 response of the CCM was determined by comparing high CO2 (SRR385610, SRR385611) against low CO2 (SRR385608, SRR385609) (Fang et al., 2012).

### Analysis of transcription factor binding sites

The coding sequence or DNA-binding domain of the gene of interest was cloned with Gibson assembly in frame with the Halo-tag into the vector pFN19A (Promega, discontinued), which contains a SP6/T7 promotor, the Halo-tag, and an Ampicillin resistance. For HY5, the codon sequence was optimized for seed plant expression and synthesized by BioCat (Heidelberg, Germany) (Table S1). Proteins were expressed using TNT® SP6 High-Yield Wheat Germ Protein Expression System (Promega, Madison, Wisconsin, United States) with 2 µg Plasmid DNA as input. The expression reaction was incubated at 30 °C for 2 h. AmpDAP-seq was performed according to Bartlett et al., (2017) with the following modifications. We used the NEBNext® End Repair Module (NEB, Ipswich, Massachusetts, United States) as the original used kit is discontinued. 70 ng of *C. reinhardtii* CC-4349 DNA library together with 1 µg UltraPure^TM^ Salmon sperm DNA solution (Invitrogen^TM^, Waltham, Massachusetts, United States) was used for the DNA-binding step as done in subsequent studies (Baumgart et al., 2021). After PCR recovery (20 cycles), the libraries were sequenced on the NextSeq 2000 using P2 reagents for 100 cycles. Expression verification of full-length fusion protein and subsequent binding to the Halo-tag beads was accomplished via Western blot analysis. The proteins were separated by 12% Tris–Glycine– SDS–PAGE and the gel was electroblotted to a PVDF membrane (0.2 µm) followed by immunodetection using Anti-HaloTag® Monoclonal Antibody (1:5000) (Promega, Madison, Wisconsin, United States) in Tris-buffered saline (TBS) including 5% milk powder, followed by a HRP-linked goat-anti-mouse IgG (1:10,000 in TBS, Agrisera AB, AS11 1772). Visualization was performed using the Pierce ECL Western blotting substrate (Thermo Fisher Scientific) and the Fusion Fx7 CCD-camera (peQLab GmbH, VWR). Reads were trimmed for adapters with Trimmomatic (Bolger et al., 2014) v0.39 and mapped with minimap2 (Li, 2018) (2.26-r1175) on the nuclear genome of *C. reinhardtii* v5.6. Reads were filtered for a mapping quality >=50 with sambamba (Tarasov et al., 2015) v0.8.0. Peaks were called with GEM (Guo et al., 2012) v.3.4 with a genome size of 107,050,806 bp and the default read distribution. Motifs were identified with MEME-Chip (Machanick and Bailey, 2011) v5.5.4 using the first 1500 peaks. Frip scores were calculated with deeptools (Ramírez et al., 2016) v3.5.1. Motifs were compared to DAP-seq motifs form *A. thaliana* with TomTom (Gupta et al., 2007) v 5.5.4. Overlaps with genomic features were determined with the R package GenomicRanges (Lawrence et al., 2013) v1.52.0. Peak summits were extended to 100 bp and assigned based of an overlap with the 200 bp upstream of the TSS, the 5’UTR and the first intron to the gene. Significant enrichment for KEGG pathways were calculated with a fisheŕs exact test.

## Results

The large wild type RNA-seq expression matrix reveals general patterns of expression and co-expression of known TFs with their target genes. All 17,741 annotated genes were expressed under at least one condition. Minimal average expression was 0.000236 transcripts per million (tpm) (Cre06.g254027) and maximal average expression was 17,263 tpm (RBCS2, a photosynthetic gene; Cre02.g120150). Rubisco small subunit 2 (RBCS2) had the highest transcript abundance under nitrogen-replete conditions at 6 h (DRX019841) and the lowest abundance under nitrogen starvation (SRX658311) with 76 tpm resulting in a dynamic range of 1,309-fold (Supplemental Table Expression Matrix). The mean abundance of all transcripts was 56 tpm and the median transcript abundance was 3.73 tpm (Figure S1). Cre06.g254027 is the gene with lowest transcript abundance and is expressed in only two experiments with very low tpm <0.2. A total of 4,943 genes are expressed in all 769 experiments (https://gitlab.ub.uni-bielefeld.de/computationalbiology/cr_grn/-/blob/main/wild-type_RNA-seq_expression_matrix.tsv.gz). The coefficient of variation over all 769 experiments ranged from 27.5 to 2,773 (Figure S2). With a coefficient of variation cut-off of 55, 273 genes can be considered housekeeping while 17,468 genes vary more widely. The median of the housekeeping gene expression is higher than the expression of non-housekeeping genes (75.5 *vs*. 5.85). The metrics indicate the data is sufficiently varied for co-expression analyses and GENIE3-based network construction.

Clustering all genes results in a heatmap indicating large diversity in expression patterns (Figure 1). Co-occurrence of PP genes (Table S4), and CCM genes (Table S4) with their known controlling TFs was expected. Nine PP genes including LHCSR1 and PSBS2 showed similar expression patterns and clustered closely together with the known TFs list here clustering away (Figure 1B). LHCRSs, PSBS1, and PSBS3 cluster distant from the other PP genes like ELIPs and PSBS2 (Figure 1B). The CCM genes and the regulators LCR1 and CrCO cluster together but are separated from the periplasmic components of the CCM and the known controlling TF CCM1 (Figure 1B). There are several conditions under which the LHCSR3s are not co-regulated with the CCM (Figure 1B). ROC75 and the CCM transporters LCIA and HLA3 are separated from the CCM cluster (Figure 1B). Taken together, the overall expression patterns in unbiased large-scale data reveal a diverse behaviour of the PP and CCM genes known to be co-expressed in some conditions (Ruiz-Sola et al., 2023; Arend et al., 2023). Co-expression with known TFs involved in these processes is limited, which indicates post-transcriptional modification, and/or non-linear, and/or combinatorial interactions.

**Figure 1:**
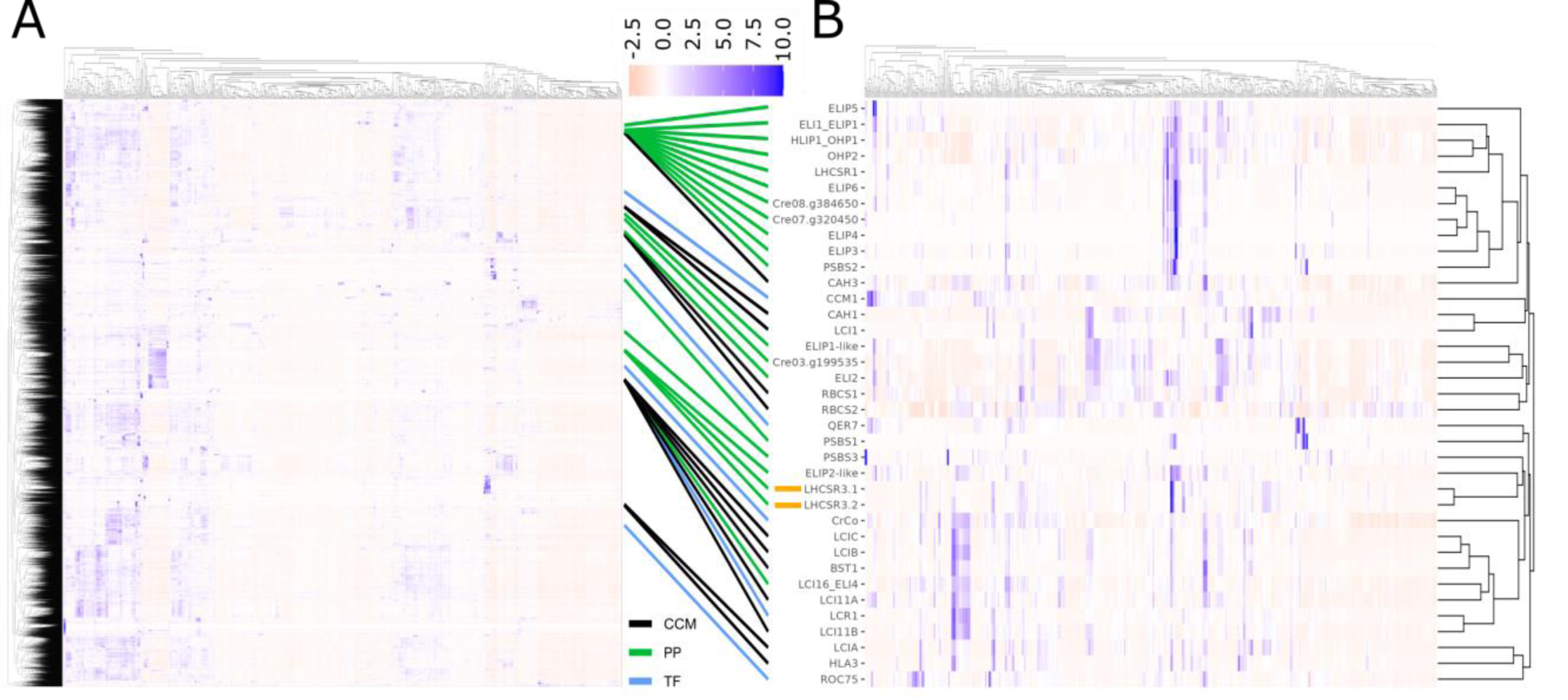
Visualization of global gene expression. **A**: Heat map of expression value of all annotated genes (z-scaled tpm (-2.5 red, white 0 and 10 blue) clustered by Euclidian distance of 769 experiments using wild-type RNA. Approximate position of genes involved in the carbon concentrating mechanism (CCM, black) or photoprotection (PP, green) and genes encoding validated transcription factors (TF, blue) is connected to B. **B**: Extracted sub-heat map of expression values of genes involved in CCM, PP, and encoding validated TFs (z-scaled tpm (-2.5 red, white 0 and 10 blue) of 769 experiments using wild-type RNA clustered by Euclidian distance (all annotated genes). LHCSR3.1 and LHCSR3.2 are marked orange.

To generate novel ranked candidate TF-target pathway associations, GENIE3-based gene regulatory networks (https://gitlab.ub.uni-bielefeld.de/computationalbiology/cr_grn/-/blob/main/GENIE3_GRN.tsv.gz) and correlation-based measures were compared for discovery potential (Figure 2). The random forest decision tree regression-based algorithm GENIE3 calculated the weighted connections of 237 TFs (Table S3) with 17,741 target genes (*C. reinhardtii* primary transcriptome v5.6), which were compared to a Pearson correlation-based analysis (Figure 2A, https://gitlab.ub.uni-bielefeld.de/computationalbiology/cr_grn/-/blob/main/person_cor_GRN.tsv.gz). Target genes for each TF were identified using a cut-off of weight (random forest-based) or correlation coefficient (correlation-based) and tested for specific and general gene ontology (GO) term enrichment as a proxy for information. Using a sliding cut-off for both networks, a similar increase in information content was observed with increasing values, but the GENIE3 network covers up to 43 unique GO term annotations for the TFs compared to 31 in the correlation-based network (Figure 2A, solid lines). Both methods reach a maximum and information content tails off with less stringent cut-offs. The GENIE3-based network was chosen for candidate discovery because of the much higher frequency of unique GO term per TF with an enrichment (Figure 2A).

**Figure 2:**
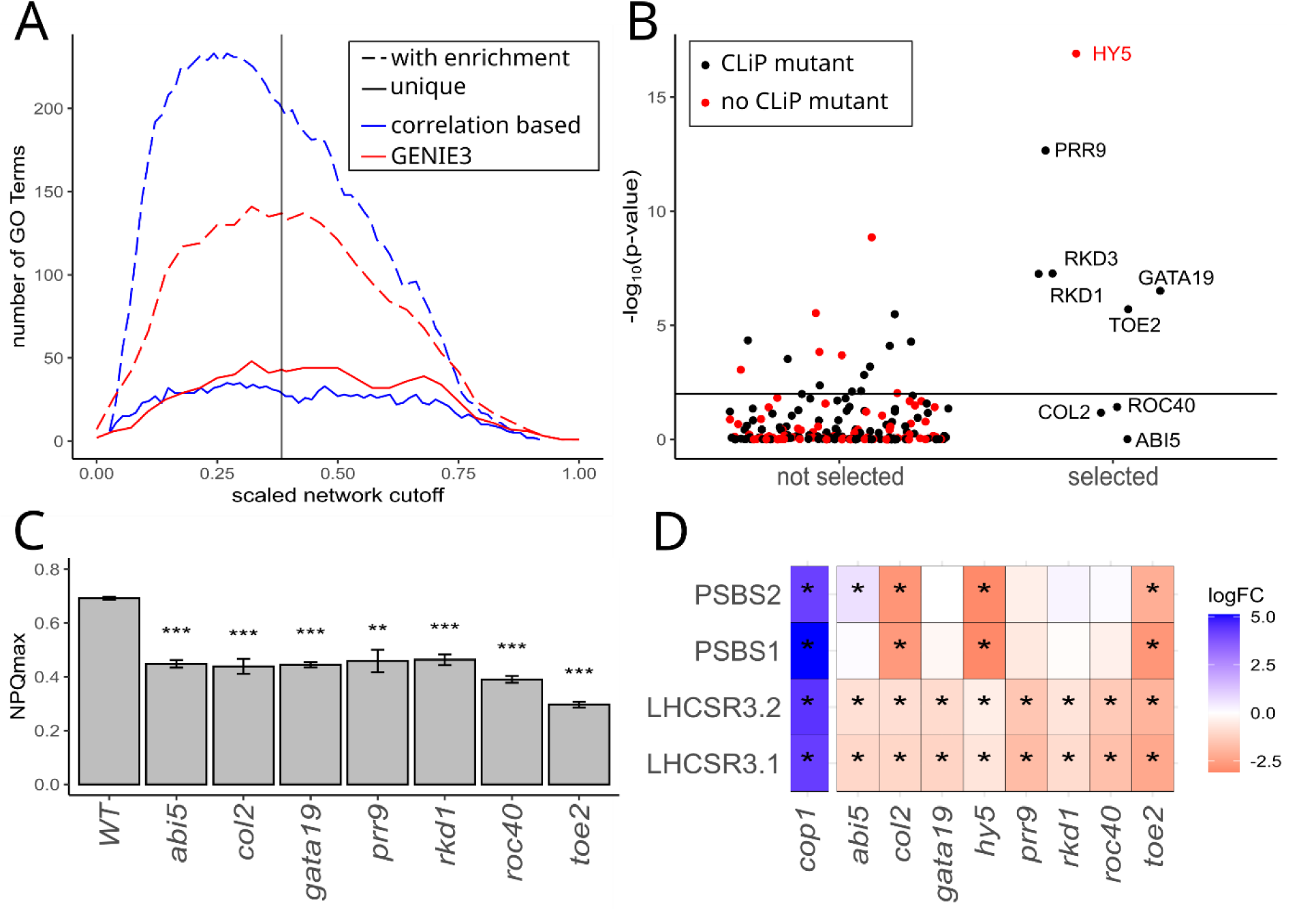
TF candidate selection and validation. **A:** Number of TFs with a GO term enrichment and number of unique GO terms with an enrichment p<1e-5 for the correlation-based network (blue) and the GENIE3 network (red) for a range of cut-offs scaled to 0 and 1. Used GENIE3 cut-off at 0.005 (scaled at 0.38) is marked with a vertical line. **B**: -log10 p-value of the Fisher exact test in the target genes of TFs from the GENIE3 network PP genes. TFs with (black) and without (red) a CliP(Li et al., 2019) mutant in the CDS selected for further investigation (right) not selected TFs (left). P<0.01 is marked with a vertical line. **C**: NPQmax of the selected, dark-acclimated CliP mutants after transfer from low light/high CO2 to high light/low CO2 conditions. Significant differences (student’s t-test) are marked with (* p<0.05, **p <0.01, ***p <0.005). Error bars indicate the standard deviation of the biological replicates. **D**: Heatmap of log2FC of PSBS1, PSBS2, LHCSR3.1, and LHCSR3.2 transcript abundances in the selected mutants and in *cop1*. Genes with reduced transcript abundances are shown in red and genes with increased transcript abundances are shown in blue. Significant differential genes (FDR<0.01, edgeR) are marked with *.

At a weight cut-off of 0.005 with custom lists (Table S5) of PP genes, 22 TFs were enriched (p<0.01). For 15 of these candidates, mutants in the CliP library are available (Figure 2B). For CCM genes, 32 TFs were enriched (p<0.01) including LCR1 and CrCO (Figure S3A). The following candidates were then selected for validation (Table 1): Cre02.g094150 (PRR9, named for homology to PSEUDO-RESPONSE REGULATOR 9 in *Arabidopsis thaliana* (Ito et al., 2007)), Cre06.g266850 (GATA19, named for homology to GATA19 (Chiang et al., 2012)), Cre03.g153050 (RKD1), Cre03.g149400 (RKD3 (Amin et al., 2023)), and Cre08.g383000 (TOE2, named for homology to TARGET OF EAT2 in *A. thaliana* (Aukerman and Sakai, 2003)) were chosen for further investigation since they score high on enrichment and there is a CliP library mutant available (Figure 2B). Three additional TFs were chosen based on their homology to *Arabidopsis* genes and the enrichment of photosynthesis genes in *C. reinhardii* (p<10e-5) (Figure S3B). These *Arabidopsis* TFs are involved in processes known to be transmitting signals to photoprotection: Cre16.g675100 (COL2) as an additional homologue of CONSTANS, which is downstream of COP1 (Liu et al., 2008), Cre05.g238250 (ABI5) as part of the ABA signal transduction (Skubacz et al., 2016), and Cre06.g275350 (ROC40) as a homologue to LHY, which is part of the morning loop in *Arabidopsis* (Mizoguchi et al., 2002). A CRISPR/Cas9 mutant was generated for Cre06.g310500 (HY5) because it had the strongest enrichment for PP. It is homologous to HY5 acting downstream of COP1 in *Arabidopsis* (Ang et al., 1998) (Table S6). RNA-seq data of *hit1-1* (*cop1)* was sampled under similar conditions (Lämmermann et al., 2020) and included in the analyses. For the CliP mutants *abi5*, *col2*, *gata19*, *prr9*, *rkd1*, and *toe2* the insertion site of the cassette at the library-predicted position was confirmed with a read length surrounding the cassette ranging from 1.1 kb to 6.0 kb (Figure S4, Table S6). No additional cassette was identified in long read data indicating no genomic rearrangements and no additional insertions are present. For the CLiP mutant *rkd3* two informative reads mapped on the cassette and the genome. One mapped from chromosome 15 over the CLiP insertion cassette into chromosome 7. The other one mapped from chromosome 15 into the cassette, then on 150 bp of the genomic sequence of RKD3 followed by the second half of the cassette. This suggested that several CLiP cassettes were inserted in the genome and a genome rearrangement happened with a likely fusion of chromosome 7 and 15. Therefore, this mutant was not analysed further (Table S6).

**Table 1:**
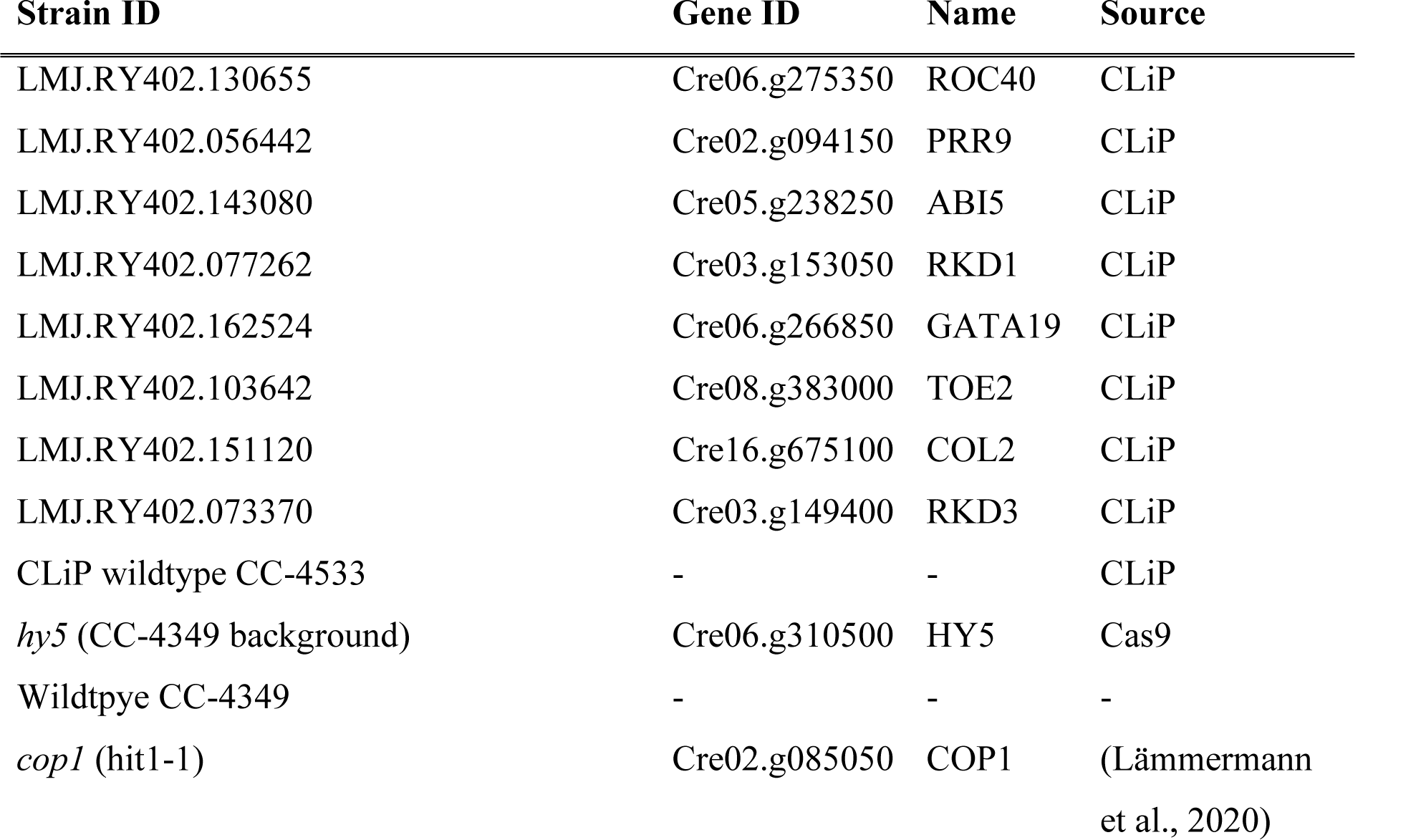
*C. reinhardtii* wildtypes and mutant lines used in this study. Presented are the strain IDs, the affected gene ID, its protein name, and the source. Mutants were either obtained from the CLiP library (Li et al., 2019) or by CRISPR/Cas9 mutagenesis.

To test if the candidate TFs play a role in photoacclimation, the verified mutant lines were tested for an NPQ phenotype and growth behaviour after a switch from low to high light conditions with limited CO2 supply. In the wild type, NPQmax was 0.70 (±.0.01 S.D.). For all mutants a significant reduction of the NPQmax was observed, with the *toe2* mutant showing the highest reduction with an NPQmax of 0.30 (±0.01 S.D.). A mild decrease in NPQmax was measured in *prr9*, *abi5*, *rdk1*, *col2,* and *gata19* and ranged between 0.44 and 0.46 (±0.01-±0.04 S.D.). In *roc40* the NPQmax was 0.39 (±0.01 S.D.). Hence, all mutants experienced a significant reduction in NPQ (Figure 2C) and in growth rate (Table S7) indicating a mutant connection to PP. We hypothesized that the changes in NPQ are a result of deregulation in the PP genes mediating or influencing NPQ. As expected from the phenotypes (Figure 2C), (Lämmermann et al., 2020), the core PP genes LHCSR3.1 and LHCSR3.2 are significantly upregulated in *cop1* and significantly downregulated in all mutants (Figure 2D). The strongest decrease in transcript abundance was observed in *toe2*, *roc40,* and *prr9*. PSBS1 and PSBS2 were significantly upregulated in the *cop1* mutant. In *hy5*, *col2,* and *toe2* both PsbS genes were significantly downregulated. In *abi5* an upregulation of PSBS2 was observed. In the remaining mutants (*prr9, roc40, rdk1,* and *gata19*) no significant change in the PsbS transcript abundances was observed. All mutants also showed dysregulation of PP genes other than LHCSRs and PsbSs (Figure S5). Especially in *col2*, *hy5,* and *toe2* most of these genes (Table S4) were downregulated. Based on the NPQ and transcriptional phenotypes, the mutated TFs are involved in PP and the population of PP-controlling TFs is larger than previously detected. The strongest reduction in the levels of all PP gene transcripts was observed in *toe2*. This coincides with the finding that this mutant was most severely affected in its NPQ capacity (Figure 2C).

We hypothesized that the comparatively large array of mutants with nine mutants under investigation with the complete transcriptome represented instead of a limited array of candidate genes tested (Redekop et al., 2022; Arend et al., 2023; Tokutsu et al., 2019) allows for the identification of photoacclimation-associated pathways (Fang et al., 2012). For better readability we distinguish between strong changes (FDR<0.01 and |log2FC|>3), moderate changes (FDR<0.01 and 3 > |log2FC| > 1) and weak changes in transcript abundance (FDR<0.01 and |log2FC| < 1). Since TFs known to be involved in PP typically also mount a transcriptional phenotype for genes involved in the CCM (Ruiz-Sola et al., 2023; Arend et al., 2023), we tested the candidate TFs for changes in CCM gene expression (Figure 3) in response to low CO2 supply (Figure 3A, red line). Under low CO2 conditions, genes of the whole CCM were upregulated (Figure 3A, Fang et al., 2012). Cytosolic carbonic anhydrase 1 (CAH1), the HCO3-transporters LCIA, HLA3, LCI11B, BST1, and the CCM components LCIC and LCIB, showed a strong, the HCO3-transporter LCI11A showed a moderate and the lumenal CAH3 showed a weak response to CO2 (Fang et al., 2012). In the TF mutants, the transcriptional response was diverse. The periplasmic components CAH1 and LCI1 had a strong positive response in *cop1.* In *abi5*, *col2*, *gata19*, *prr9*, *rkd1*, *roc40 and toe2* a strong downregulation was observed with the exception of *hy5* where CAH1 was weakly downregulated (Figure 3A). HLA3 was downregulated in all mutants except for *hy5* with an unchanged expression. In *cop1* and *toe2* this downregulation was moderate and in *abi5*, *col2 gata19*, *prr9*, *rkd1,* and *roc40* it was weak. The plastidic transporter LCIA was again upregulated in *cop1* and significantly downregulated in all mutants except *hy5*. The downregulation of LCIA was strong in *prr9*, moderate in *toe2*, *col2*, *rkd1,* and *roc40* and weak in *abi5* and *gata19* (Figure 3A). The thylakoid transporter BST1 was strongly upregulated in *cop1*, moderately downregulated in *prr9* and *toe2*, and weakly downregulated in *rkd1*, and *roc40*.

**Figure 3:**
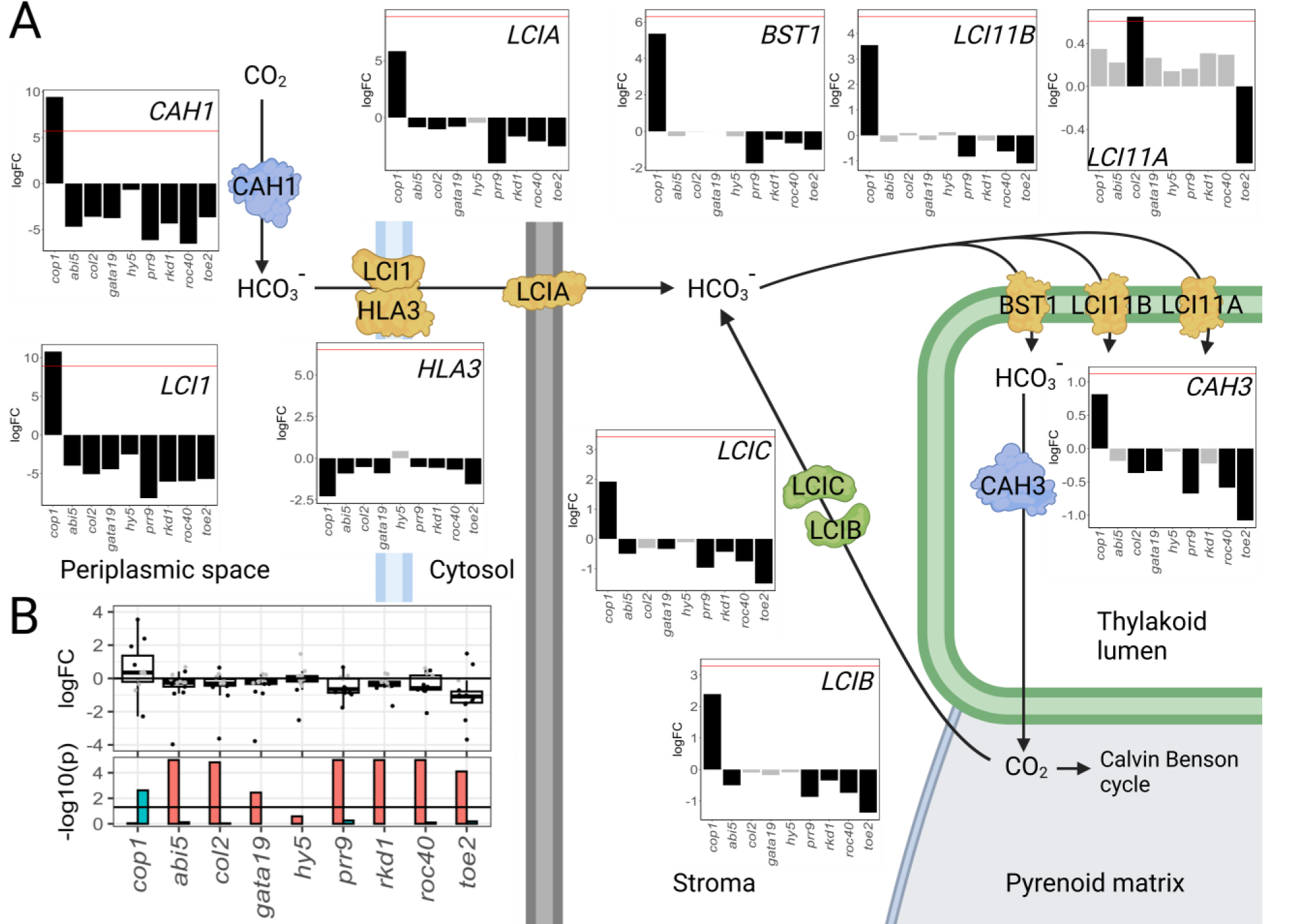
Effects of mutated TFs on transcript abundances of CCM genes. **A:** CCM pathway in *C. reinhardtii* with bar plots of the log2-fold change of the mutants *abi5*, *col2*, *cop1*, *gata19*, *prr9*, *rkd1*, *roc40* and *toe2*. Significantly different genes (FDR<0.01) are shown as black bars, not significantly differential genes (FDR>=0.01) as grey bars. The red line indicates the physiological response of *C. reinhardtii* under low CO2 conditions (400 ppm CO2 in air). **B:** Top panel: Boxplot and jitter plot of individual genes of log2-fold change of CCM genes. Significantly different genes (FDR<0.01) are shown in black, not significantly differential genes (FDR>=0.01) are marked grey. In the bottom panel is a bar plot of the –log10 p-value capped at 1e-5 of the gene list enrichment of the significantly upregulated genes (green) and significantly downregulated genes (red). **Created with** BioRender.com

The thylakoid transporter LCI11B was strongly upregulated in *cop1,* moderately downregulated in *toe2* and weakly downregulated in *prr9* and *roc40*. LCI11A showed the weakest induction under low CO2 conditions and different dysregulation in the mutants compared to the other transporters (Figure 3A). The thylakoid lumen carbonic anhydrase CAH3 was weakly upregulated in *cop1* and weakly downregulated in *col2*, *gata19*, *prr9,* and *roc40* and moderately downregulated in *toe2*. LCIB and LCIC showed a similar pattern (Figure 3A). Both were moderately upregulated in the *cop1* mutant and moderately downregulated in *toe2*. A weak downregulation was observed in both for *abi4*, *prr9*, *rkd1*, and *roc40*. For *gata19* and *col2* the downregulation was only observed in LCIC. Overall, except for LCI11A, in *cop1* an upregulation of the CCM was observed whereas in all other mutants except *hy5* a downregulation was observed. The upregulation in *cop1* was on average higher than the downregulation in the other mutants. The strongest downregulation was observed in *prr9* (mean: -2.4, median: -0.9) followed by *toe2* (mean: -2.0, median: -1.4) and *roc40* (mean: -1.8, median: -0.7). When the CCM is defined as the genes, which encode the pyrenoid (Table S4), all mutants except *hy5* and *cop1* were significantly enriched in downregulated genes (Figure 3B). The *hy5* mutant was not enriched at all, while *cop1* was enriched in CCM genes among upregulated genes (Figure 3B). We conclude that in the majority of mutants in which PP is deregulated (Figure 2D), the CCM is also altered in expression, however, the degree of change varies and is absent for *hy5* (Figure 3 A, B). Since the mutants of PP TFs are altered in expression for PP and for CCM genes and thus involved in the photoacclimation regulon, we hypothesized that they can be applied to identify other pathways that are potentially associated with photoacclimation.

The mutants provided a unique opportunity to examine photosynthetic modules potentially linked to increasing electron sinks or reduction in harvested photons (Figure 4, Figure S6). In the *toe2* mutant, genes encoding enzymes in the CBBc cycle were upregulated, as were most of the LHCs with the exceptions of LHCB7 and LHCBM9 (Grewe et al., 2014) (Figure S6E). All nuclear-encoded PSI genes were upregulated and all nuclear-encoded PSII genes with the exceptions of PSBP1-3 and PSB28 were upregulated (Figure S6E). ATPase was unchanged (Figure S6E). Similar patterns were observed in all mutants (summarized in Figure 4D-I, Figure 4A-C, Figure S6) with some exceptions. The LHCs were significantly upregulated in *abi5, col2*, *gata19, prr9, rkd1, roc40,* and *toe2*. The upregulation was in a similar strength with a median log2-FC between 1.1 and 0.8 except for *prr9* where a weaker upregulation of log2-FC 0.42 was observed (Figure 4D). In *cop1* and *hy5* the opposite behaviour of these genes was observed with a median log2-FC of -0.45 in *hy5* and -1.3 in *cop1*. For the nuclear genes encoding the photosystems, significant upregulation was observed in *abi5*, *gata19, rkd1*, *roc40,* and *toe2* (Figure 3E). In *cop1* a significant downregulation of the photosystems was observed. The CBB cycle was significantly upregulated in *abi5*, *gata19*, *prr9*, *rkd1*, *roc40,* and *toe2*, but was unaltered in *col2*, *cop1,* and *hy5* (Figure 4F). Overall, LHCs, PSs, and CBBc appear co-regulated (Figure 4D-F) and regulated in the opposite direction to the PP and CCM genes (Figures 2D, 3A, B). Taken together, the TFs controlling photoacclimation also control photosynthetic modules but changes are not coordinated. In stark contrast, photorespiration was significantly downregulated in many mutants (Figure 4G). In *abi5*, *gata19, prr9*, *rkd1*, *roc40,* and *toe2* the pathway was downregulated with the strongest effect in *toe2*, *prr9,* and *roc40* and no changes in *col2, cop1,* and *hy5* (Figure 4G).

**Figure 4:**
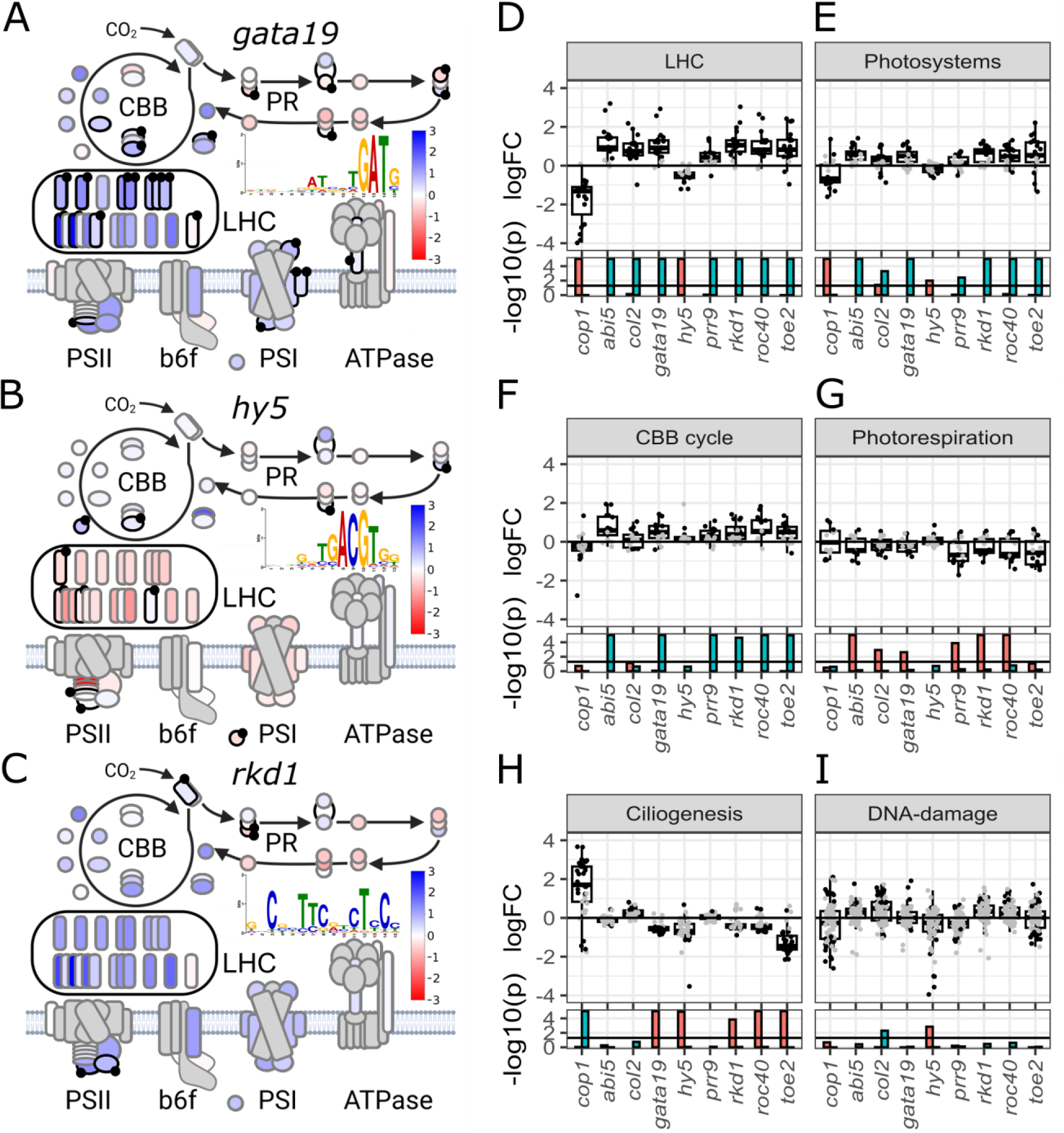
Effects of mutated TFs on transcript abundances in selected KEGG pathways. **A-C**: Photosynthesis pathway heatmap of *gata19*, *hy5*, and *rkd1* of the log2-FC (-3 red, 0 white, 3 blue) of the CBB, PR, LHC, PSII, b6f, PSI and ATPase. Plastid encoded genes are filled in grey. The binding motif of the TF was determined with DAP-seq is depicted in the middle. Bound genes determined with DAP-seq have a black boarder and are marked with a black Other mutants and significance of the log2-FC is depicted in **D-I** in addition with the pathways ciliogenesis and DNA-damage: Top panel boxplot of the log2-FC and jitter of individual genes of the mutants *abi5*, *col2*, *cop1*, *gata19*, *hy5*, *prr9*, *rkd1*, *roc40* and *toe2*. Significantly differential genes (FDR<0.01) are black, other genes are grey. The bottom panel bar plot of the -log10 p-value capped at 1e-5 of the gene list enrichment of the significantly upregulated genes (green) and significantly downregulated genes (red). **Created with** BioRender.com

Unbiased tests for KEGG pathways showed 45 out of 238 modules significantly changed in the mutants tested (summarized in Table S9, Figure S8). In addition, changes in ciliogenesis were tested based on the changes in swimming behaviour upon ABA treatment (Al-Hijab et al., 2019) and light stress (Erickson et al., 2015). Changes in ciliogenesis coincided with those observed with PP and the CCM in some of the mutants (Figure 2D, Figure 3B, Figure 4H). In *cop1* overall an upregulation with a median log2-FC of 1.70 of these genes was observed. In *col2* a weak upregulation (log2-FC: 0.24) of ciliogenesis was observed. A significant downregulation of these genes was observed in *gata19*, *hy5*, *rkd1*, *roc40,* and *toe2* with median log2-FC between -0.4 and -0.7 (Figure 4H). The DNA-damage and DNA-repair pathway was significantly upregulated in *col2*. In the *hy5* mutant 13 DNA-damage-related genes were significantly downregulated with a median log2-FC of -1.66. Especially Cre06.g278151, RAD54B and RAD51 showed the strongest downregulation (log2-FC<-3). In the *abi5*, *rkd1,* and *roc30* DNA-damage-related genes were overall upregulated. On average, the mutants *cop1*, *gata19,* and *toe2* were not altered in the transcript abundances in this pathway (Figure 4I). In *toe2* and *col2* carotenoid biosynthesis was significantly downregulated with a median log2-FC of -0.94 (*toe2*) and -0.6 (*col2*) (Table S9, Figure S8). The TF mutants, which were affected in NPQ and PP gene expression demonstrate that their target gene regulon extends beyond the acclimation genes and includes LHCs, PSs, CBBc, PR, ciliogenesis, and DNA damage and repair pathways (Figure 4), among others (Table S9, Figure S8).

To distinguish between direct and indirect actions of TFs on target genes and to test whether the TFs bind each other’s promoters, we attempted DAP-seq for all TFs under analysis and succeeded for the full-length protein of HY5 and the DNA-binding domain of RKD1 and GATA19. By western blotting we confirmed the expression of the proteins (Figure S7A). RKD1 bound regions with the motif (CNNTTCNNCTNC) (Figure 4C), which was most similar to the motif of At-RKD2 DAP-seq data (q<1.79e-04). The HY5 motif was the G-box (ACGTG) (Figure 4B), which overlapped strongest with bzip TFs (At-bzip50, q<1.95e-04). The GATA19 motif was ATNTGATG (Figure 4A), which had the most significant overlap with the motif of GATA19 (q<9.18e-08). Hence, binding motifs are strongly conserved for these three TFs between the seed plant *A. thaliana* and the microalga *C. reinhardtii*, between which lies an evolutionary distance of 675-850 million years (Harris et al., 2022). By assigning target genes based on peaks overlapping with 200 bp upstream of the TSS, the 5’ UTR and first intron, this resulted in 3,355 bound genes for GATA19, 1,576 for HY5 and 1,161 for RKD1 (Table S11). All TFs shared 54 target genes, HY5 and GATA19 shared 451 target genes, RKD1 and GATA19 shared 324 target genes, and HY5 and RKD1 shared 148 target genes. Most genes were specific for each of the tested TFs (Figure S7B). Enrichment of pathways among targets (p<0.1) for GATA19 identified a enrichment common to bound genes and deregulated genes (Kanehisa et al., 2017) in photosynthesis antenna proteins, “N-glycan biosynthesis”, cytoskeleton proteins, and amino acid related enzymes (Figure S8, Table S12). For RKD1 a common enrichment was detected for ribosome biogenesis, cytoskeleton proteins, motor proteins, N-Glycan biosynthesis, protein kinases, and starch and sucrose metabolism (Figure S8, Table S12). In HY5 the bound and deregulated genes had a pathway enrichment in “protein digestion and absorption” and ubiquitin systems (Figure S8, Table S12). Direct targets among known photoacclimation genes, photoperception genes and TFs were manually analysed. HY5 bound to aCRY, pCRY, and CRY-DASH2 and these genes were not bound by RKD1 and GATA19 (Table S11). RKD1 bound to the CCM regulator LCR1 and the photosynthesis regulator CrCo. HY5 bound QER7. The carbonic anhydrase CAH1 was bound by RKD1 and GATA19 (TableS4, Table S11). Out of the PP genes GATA19 bound ELIP5, PSBS3, ELI2 and Cre03.g199535. HY5 bound ELIP5 and Cre07.g320450. RKD1 bound to ELIP1 (Table S4, Table S11). GATA19 bound several PSII related genes PSBP6, PSAE, PSAH, PSAK, and PSAN. HY5 bound to PSBQ, PSBP4, and PCY1. RKD1 bound PSBQ and PSBR. The three TFs also bound CBBc genes. GATA19 bound to four CBBc genes (GAP1, TPIC1, FBA1, FAB3). HY5 bound to FAB3 and FBP1. RKD1 bound to RBCS1. For PR RKD1 and HY5 bound two genes (RKD1: Cre03.g168700, Cre06.g271400; HY5: SHMT2, Cre06.g278148). GATA19 bound to five PR genes (Cre03.g168700, GYX1 SHMT2, SHMT1, Cre06.g278148) (Figure 4A-C, Table S11). Taken together the analyses for direct or indirect effects demonstrated that the three TFs analysed acted both directly on their target genes, and indirectly via other TFs and via photoperception genes.

## Discussion

Analyses of the mutants for photoacclimation-controlling TFs demonstrate the complex interaction and action of TFs in the photoacclimation gene regulation (Figure 5) that reflects the complexity of eukaryotic TF-based gene regulation to both input pathways and target genes and modules. The initial analysis of the co-expression matrix of CCM genes, PP genes and their known regulators suggested that regulatory processes are complex and potentially combinatorial since co-expression of the known genes among available datasets is limited (Figure 1B). Indeed, the RF approach of candidate identification (Figure 2A) and validation (Figures 2B, C, and D) identified nine additional TFs with a role in PP that contribute to the varying modules of PP in a wider sense. Validation via long read genome resequencing demonstrated that at least some of the CLiP library mutants suffer from large-scale re-arrangements potentially induced by multiple insertions of the mutation cassette. One mutant, the candidate RKD3 mutant, had to be excluded from the analysis based on the detection of a genome rearrangement in the long reads (Figure S4, Table S6). Genome rearrangements in insertion mutants were previously observed in up to 10% of T-DNA insertion mutants in *A. thaliana* (Pucker et al., 2021). The majority of mutants however do not show evidence of multiple insertions or genome rearrangements (Figure S4, Table S6).

**Figure 5:**
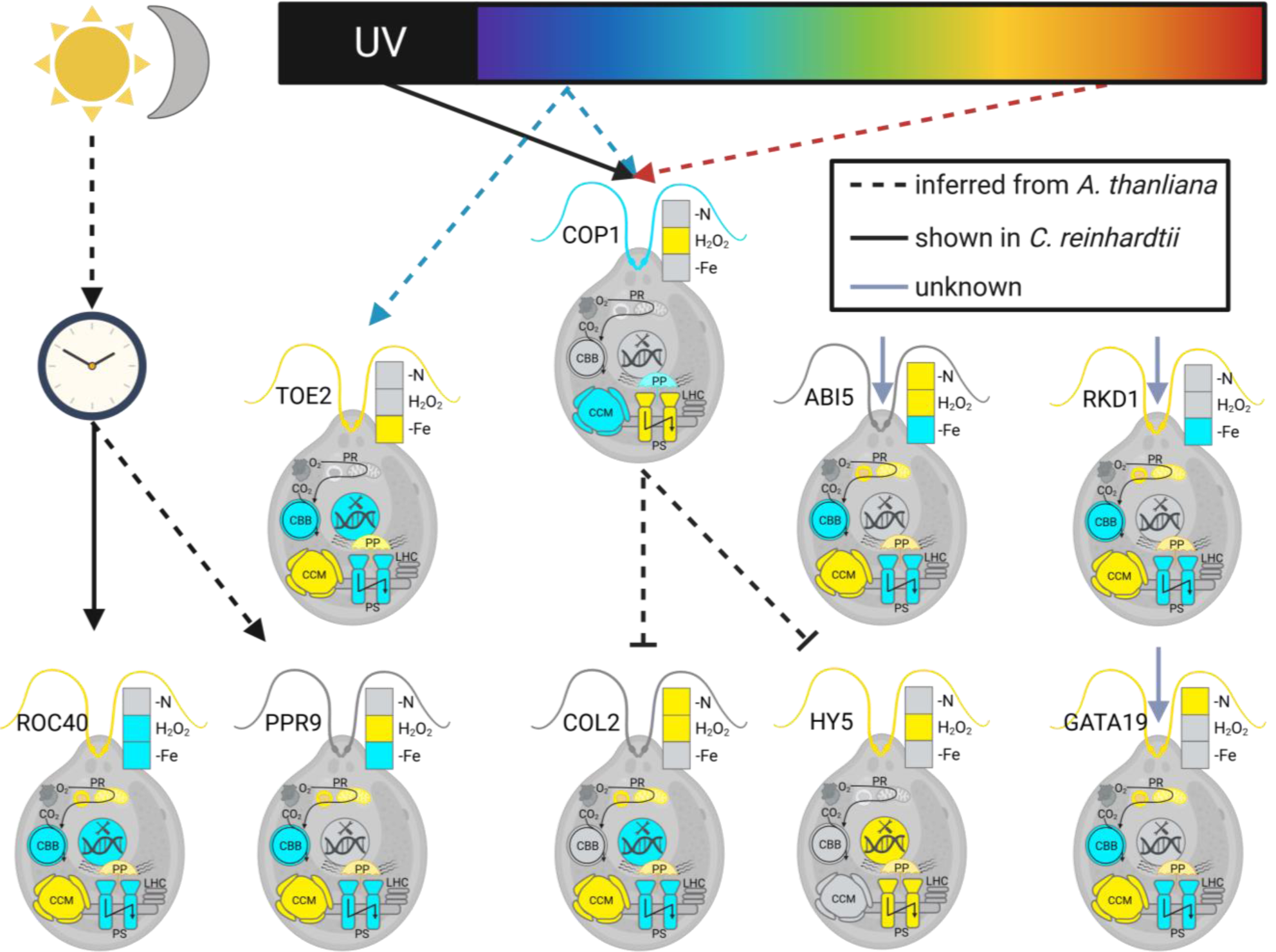
Model of the response of the TFs to various stressor inputs, such as Nitrogen deficiency (-N) Oxidative stress (H2O2) and Iron deficiency (-Fe) and outputs of TFs on the processes Ciliogenesis, Photorespiration (PR) (Table S13), Calvin Benzon Benson Bassham cycle (CBBc), carbon concentrating mechanism (CCM), Photosystems (PS), light harvesting complexes (LHC), photoprotection (PP), DNA-repair (yellow positive regulation and blue negative regulation). Solid black arrows mark connections shown in *C. reinhardtii*, dashed arrows mark connections inferred from *A. thaliana* and solid grey arrows mark currently unknown inputs. **Created with** BioRender.com

For the remaining TF mutants, the RNA-seq data of *cop1*, *hy5*, *col2*, *toe2*, *roc40*, *prr9*, *gata19*, *rkd1,* and *abi5* suggested a model in which enabling high light acclimation extends beyond PP genes *sensu strictu* and the CCM (summarized in Figure 5). In seven mutants, photorespiration is affected and co-downregulated with PP genes as was observed previously for the *ccm1* mutant (Fang et al., 2012). In seed plants, PR has been suggested as a major electron sink for photoprotection (Bauwe et al., 2012; Eisenhut et al., 2017) although C4 plants survive high light with little to no PR (Mallmann et al., 2014). The PR pathway consumes 2 NADPH and 3.5 ATP when the oxygenation product of Rubisco, phosphoglycolate, is recycled to the CO2 acceptor ribulose-1,5-bisphosphate (Walker et al., 2016). Hence it is an efficient way to dissipate excess energy without damaging the cell when energy equivalent and reductant consumption by the CBBc is low (Villalobos-González et al., 2022). This photorespiratory energy valve is probably also active in algae based on its co-regulation with photoacclimation genes. The acclimation in the architecture and the amount of light-harvesting and photosystem complexes to the amount of available light known from seed plants (Eisenhut et al., 2013) is only partially observed in the alga. In eight mutants, LHC and PS genes are inversely regulated compared to PP genes (Figure 3, Figure 5), but no changes in PSI to PSII ratio (Bailey et al., 2001) or a decoupling of LHCs and PS from the CBBc can be observed (Halpape et al., 2023). The observed general reduction of expression for LHCs and PS may ultimately lead to a reduction in light-harvesting capacity and therefore to reduced pressure on PP. The energy-demanding process of swimming with their ciliae (Brokaw et al., 1982) may contribute two-fold to photoacclimation *sensu lato*. The process itself consumes energy in the form of ATP (Burgess et al., 2003) and it may be used to alter the position of the alga within the water column and thereby adjust the amount of light and light quality (Hegemann and Berthold, 2009). Finally, light stress can induce ROS, which can lead to DNA-damage (Apel and Hirt, 2004) and, consequentially, DNA repair is altered in *hy5* and *col2* (Figure 4, Figure 5). The expression changes of photosynthetic modules and other pathways thus contribute to acclimation to excess light but act slightly differently compared to the seed plant *A. thaliana*.

The parallel analyses of nine TFs and COP1 in mutants, their partially known linkage based on data from *C. reinhardtii* (Han et al., 2020) and their homology to proteins with known functions in the seed plant *A. thaliana* allow the construction of a schematic model (Figure 5). While the transfer of data from a seed plant to an alga may seem like a reach, data gathered about HY5 suggests a large degree of conservation. As found in *A. thaliana* direct targets of HY5 in *C. reinhardii* are genes related to the E3 ligase complex (Burko et al., 2020). In addition, DAP-seq analysis showed that in *C. reinhardtii* aCRY, pCRY, and CRY-DASH2 are direct targets of HY5 (Table S11). As in *A. thaliana* this primes *C. reinhardtii* for a rapid response upon light exposure (Burko et al., 2020). In a *C. reinhardtii* regulatory model, the eight TFs and COP1 can be placed into a putative light regulon and an unknown category. Re-analysis of published RNA-seq data also demonstrates that these TFs respond to other inputs, such as nitrogen status, iron status, and hydrogen peroxide (Figure 5). In the light regulon, a (UV-) light signal is likely transmitted via COP1 (Favory et al., 2009; Allorent et al., 2016) to HY5 and COL2 based on the homologous role of the downstream TFs in *A. thaliana* (Gangappa and Botto, 2014) (Figure 5). In *A. thaliana*, COP1 acts downstream of both red and blue light (Ponnu and Hoecker, 2021), raising the question of whether the same is true for *C. reinhardtii* COP1. PP genes are inversely regulated in *crcop1* compared to *hy5* and *col2* (Figure 5), which indicates a similar role of CrCOP1 to At-COP1’s inhibitory role in *A. thaliana* (Ponnu and Hoecker, 2021). The RNA-seq identified downstream targets HY5 and COL2 diversify responses of CCM, LHCs, PS, and PR (Figure 5) allowing a tuning of the response based on other potential interactors and input pathways. Data in *A. thaliana* suggests multiple modulators of HY5 and the BBX-type COL2 TF (Gangappa and Botto, 2016; Job et al., 2018; Burko et al., 2020; Serrano-Bueno et al., 2020; Zhang et al., 2022; Cao et al., 2023) including a proposed interaction of HY5 with BBX-type TFs, such as COL2 (Gangappa et al., 2013). TOE2 is tentatively placed downstream of light since the homologue in *A. thaliana* transmits the blue light signal perceived via CRY (Ahmad et al., 1998). At-TOE2 interacts with At-CRY1/2 and inhibits At-COL1 and CO and antagonizes the flowering time in *A. thaliana* (Zhang et al., 2015; Du et al., 2020). In the *acry* mutant (Beel et al., 2012) carotenoid biosynthesis is deregulated as it is in *toe2* (Table S10, Figure S8) making it tempting to place aCRY upstream of TOE2. TOE2 has the strongest effect of all mutants examined. In the *toe2* mutant most genes were differential (Table S9). It shows overall the strongest negative effects of all the examined mutants and it showed the strongest NPQ phenotype (Figure 2C). The other TFs under investigation except ROC40 are positively regulated by TOE2 (Figure S9). Two clock genes, ROC40 (known as LHY in *A. thaliana*) and PRR9 also show deregulation of PP (Figure 5). Light entrains the clock (McClung, 2006) and clock mutants have defects in leave movement and flowering time control in *A. thaliana* (Mizoguchi et al., 2002). It is at first glance surprising that both the morning and the evening circuit mutants show similar responses (Figure 5). LHCSR genes peak twice during the day, in the morning and the afternoon (Strenkert et al., 2019) (Figure S5), which may point to both circuits contributing to their expression. In addition, the clock period is altered in *roc40* (Matsuo et al., 2008) and potentially altered in *prr9* if its role is similar to that in Arabidopsis (Nakamichi et al., 2005), which may shift typical gene expression peaks (Strenkert et al., 2019). The transcriptional and NPQ phenotypes of the mutants of *cop1*, *hy5*, *col2*, *toe2*, *roc40*, and *prr9* strengthen the evolutionary link of PP and flowering (Tokutsu et al., 2019). Light protective circuits in *C. reinhardtii,* which necessarily sense, and/or transmit, and/or compute light signals have evolved into circuits, which by virtue of their ability to process light regulate flowering that depends on time of year and therefore day length in seed plants. The four additional TFs chosen based on their enrichment of targets in PP (Figure 2B) and based on homology with TFs in *A. thaliana* suggest links to known signalling pathways based on their homology to *A. thaliana*. GATA19 is homologous to GATA sub-family II TFs in *A. thaliana,* which act downstream of HY5 (Manfield et al., 2007; Chattopadhyay et al.; Zhang et al., 2022) and influence chlorophyll content, photomorphogenesis and flowering. At-GATA19 is involved in the regulation of nitrogen deficiency (Shen et al., 2022) (Figure 5) and cytokinin (Chiang et al., 2012). In *C. reinhardtii* the nitrogen responsive GATA19 (Figure 5) directly binds photosynthesis genes indicating it links the nitrogen towards photosynthesis. The RWP-RK TFs are of the RKD(B) subfamily, which is expanded in *C. reinhardtii* compared to seed plants (Chardin et al., 2014). Members of these families have diverse known roles, nitrogen responses and gametophyte development in seed plants and in gametogenesis in *C. reinhardtii* (Chardin et al., 2014; Lin and Goodenough, 2007). The RKD1 is transcriptionally reduced under iron deficiency and does not show a response under nitrogen deficiency, and H2O2 stress (Figure 5). The varied responses to cues other than light and CO2 suggest that these TFs serve to integrate the PP response with other stress responses (Figure 5). The bound genes of RKD1 only included a minority of genes involved in CCM, PP, and PS. Enrichments were in cytoskeleton, motor proteins, ribosome biogenesis, N-Glycan biosynthesis, and starch and sucrose metabolism. Thus, the observed phenotype is likely a result of secondary effects. For example, RKD1 may alter the expression of regulators involved in these processes or it changes expression of the bound key genes e.g., PSBR or CAH1 catalysing the first step of the CCM, which in turn may be sufficient to alter functions of the pathways leading to secondary effects. The analysis of the Cr-ABI5 mutant demonstrates the limits of information transfer from seed plants to algae. It was originally chosen because At-ABI5 mediates abiotic stress responses in seed plants through the ABA-signalling pathway. In *C. reinhardtii*, ABA modulates HCO3-uptake and relative position of *C. reinhardtii* in the water column (Al-Hijab et al., 2019). While the mutant displays weak altered PP responses (Figure 2C), the expression of at least one PsbS gene is opposite expectations (Figure 2D). Of all mutants analysed, its transcriptional connection to PP (Figure 2D) is the weakest, which reflects its low position on the target gene enrichment analysis (Figure 2B).

The analysed TFs constitute a large group of TFs involved in photoacclimation that acts together with CCM1/CIA5, LCR1, and QER7. RNA-seq analyses of nine regulator mutants show that COP1 and the TFs influence each other in expression (Figure S9) and form a network rather than a hierarchical system. While single TFs, such as TOE2 may have an outsized impact (Figure 2C-D, Figure3, Figure 4, Figure S5, Figure S9), they are still part of a redundant network that mellows the effect of loss (i.e. Figure 2D, Figure 3). A signal itself, such as decreased CO2 leads to large amplitudes of changes in CCM transcripts (Figure 3A) compared to the signals induced by each TF mutant. Each TF mutant itself shows statistically significant, detectable changes in single genes (Figure 3A) and in the group (Figure 3B). However, these changes are smaller compared to the transcriptional changes observed upon the signal itself. Similarly, the changes in the PP genes LHCSRs are modest with downregulation of up to 2.5-fold in each mutant with concomitant modest changes in NPQ (Figure 2C). NPQ phenotype and transcript reduction in LHCSRs appear correlated in *toe2* with the strongest changes, but not in the other mutants (Figure 2C, D). NPQ is indeed modulated by other processes (Redekop et al., 2022), which may break the absolute link between NPQ phenotype and RNA-seq data. The global RNA-seq results point to shared regulation of expression between more than ten different TFs acting in a robust network based on partial redundancy. The very large, up to 1,370-fold changes in gene expression detected in the expression matrix are thus likely also the result of multiple TFs acting jointly rather than a single regulator. The global RNA-seq analyses of multiple TFs acting redundantly in the same regulon reflect the complexity of eukaryotic gene regulation where dozens of TFs bind the same promoter region (Franco-Zorrilla et al., 2014; Brkljacic and Grotewold, 2017; O’Malley et al., 2016; Zenker et al., 2023) and act in concert to influence gene expression (Brkljacic and Grotewold, 2017).

## Data availability

Raw sequencing data of RNA, long read DNA-seq of the CLiP mutants and DAP-seq of HY5, RKD1 and GATA19 is available under the following Bioproject. https://dataview.ncbi.nlm.nih.gov/object/PRJNA1020056?reviewer=i385dk6uclut8o7pu8hd638dfg

The GRN and expression matrix, DAP-seq peaks, and motifs is available in this data publication: https://gitlab.ub.uni-bielefeld.de/computationalbiology/cr_grn

## Acknowledgements

The authors gratefully acknowledge Tobias Busche and Yvonne Kutter for their valuable support with genome sequencing and the Chlamydomonas Mutant Library Group at Princeton University, the Carnegie Institution for Science, and the Chlamydomonas Resource Center at the University of Minnesota for providing the indexed Chlamydomonas insertional mutants. We are grateful to the Center for Biotechnology (CeBiTec) at Bielefeld University for access to the Technology Platforms.

This work was supported by the BMBF-funded de.NBI Cloud within the German Network for Bioinformatics Infrastructure (de.NBI) (031A532B, 031A533A, 031A533B, 031A534A, 031A535A, 031A537A, 031A537B, 031A537C, 031A537D, 031A538A).

Figure 3, Figure 4A-C and Figure 5 were created with BioRender.com.

## Author Contributions

D.W. designed and carried out computational analyses (DEG analysis, confirmation of CLiP insertion mutants, DAP-seq data) and designed and carried out DAP-seq experiments, interpreted data and co-wrote the paper.

F.J.K. carried out computational and wet lab experiments and interpreted data and edited the paper.

L.J.K carried out the nanopore sequencing and edited the paper.

E.J. provided the Cas9 insertion mutant of HY5 and edited the paper.

P.V. planned and carried out sequencing.

L.W. interpreted data and edited the paper.

E. M. interpreted data and co-wrote the manuscript.

O.K. interpreted data and edited the paper.

O.B.K. designed and carried out wet lab experiments, interpreted data and co-wrote the paper.

A.B. conceived the initial idea and the study, interpreted data and co-wrote the paper.

## Supplemental materials

Supplemental

Supplemental figures 1-9 and Supplemental tables 1-13 are available as separate files. Supplemental data is available under https://gitlab.ub.uni-bielefeld.de/computationalbiology/cr_grn and will be made available published as a data publicatoin.

