## Supplemental Figures for "Multiple transcription factors mediate acclimation of Chlamydomonas to light stress"

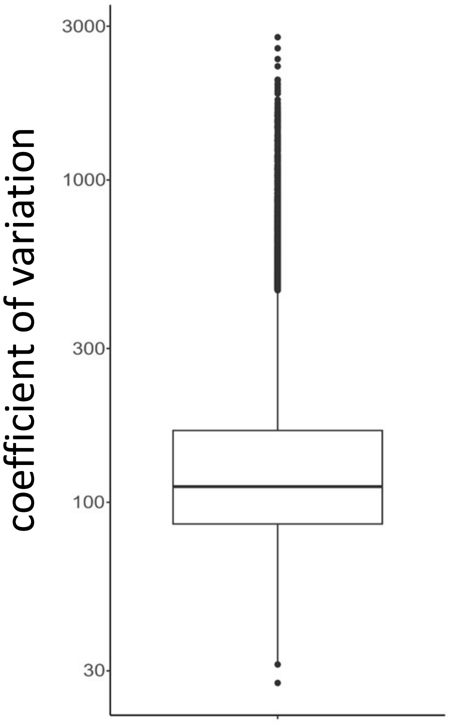


*Figure S1: Boxplot of coefficient of variation of all expressed genes in 769 C. reinhardtii wild type experiments*

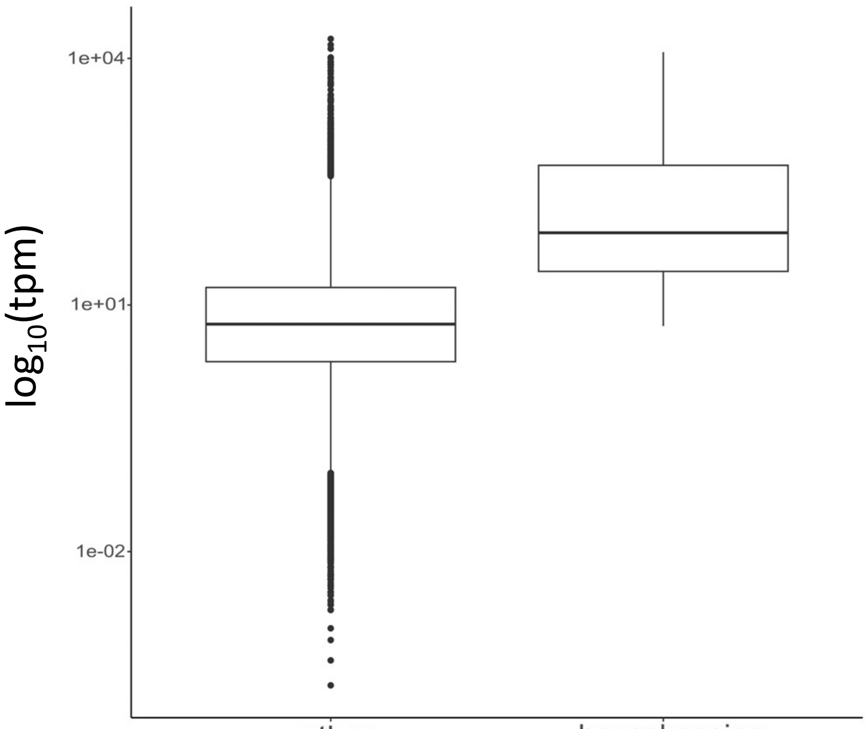


*Figure S2: Boxplot of the median $log_{10}$(tpm) of housekeeping genes (cv<55) and other genes (cv>=55).*

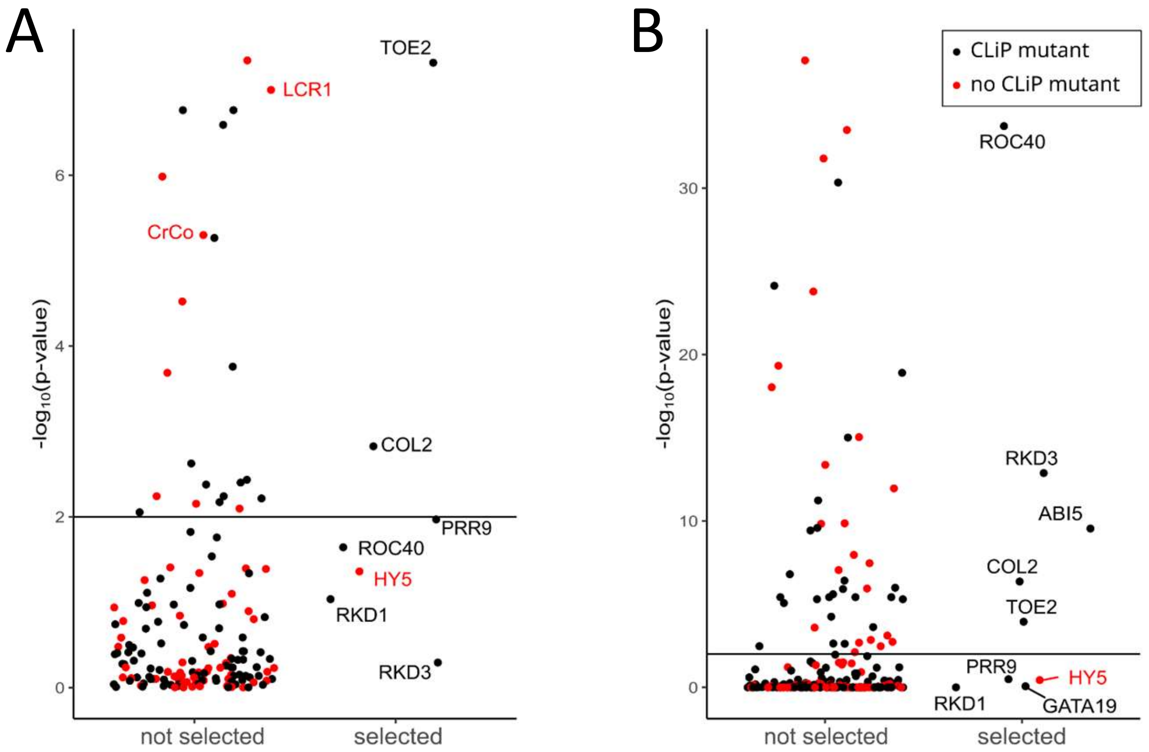


*Figure S3: -log10 p-value of the Fisher exact test in the target genes of transcription factors from the GENIE3 network of the CCM (A) and photosynthesis genes (B). Transcription factors with (black) and without (red) a CLiP45 mutant in the CDS selected for further investigation (right) not selected transcription factors (left).*

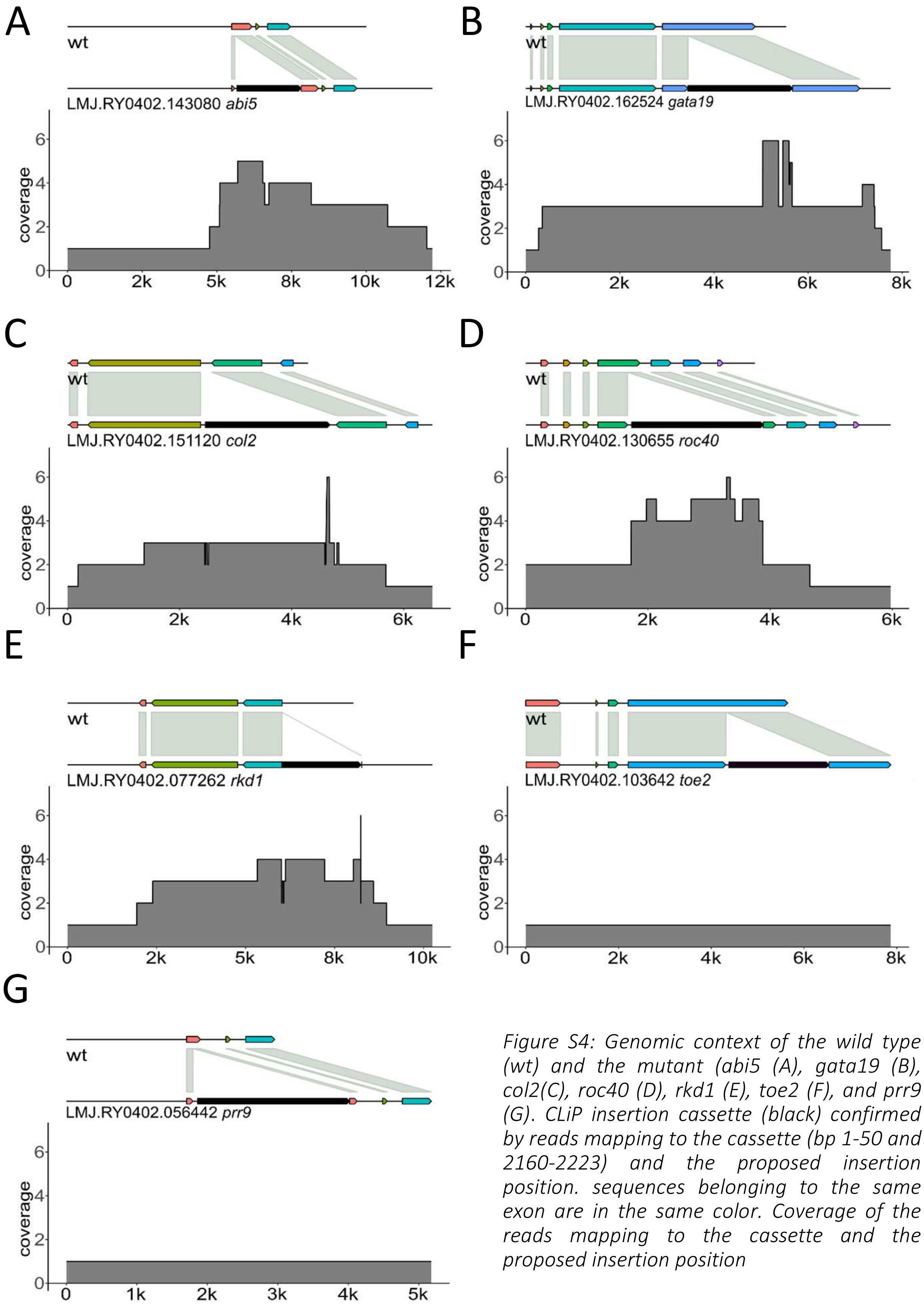


*Figure S4: Genomic context of the wild type (wt) and the mutant (abi5 (A), gata19 (B), col2(C), roc40 (D), rkd1 (E), toe2 (F), and prr9 (G). CLiP insertion cassette (black) confirmed by reads mapping to the cassette (bp 1-50 and 2160-2223) and the proposed insertion position. sequences belonging to the same exon are in the same color. Coverage of the reads mapping to the cassette and the proposed insertion position*

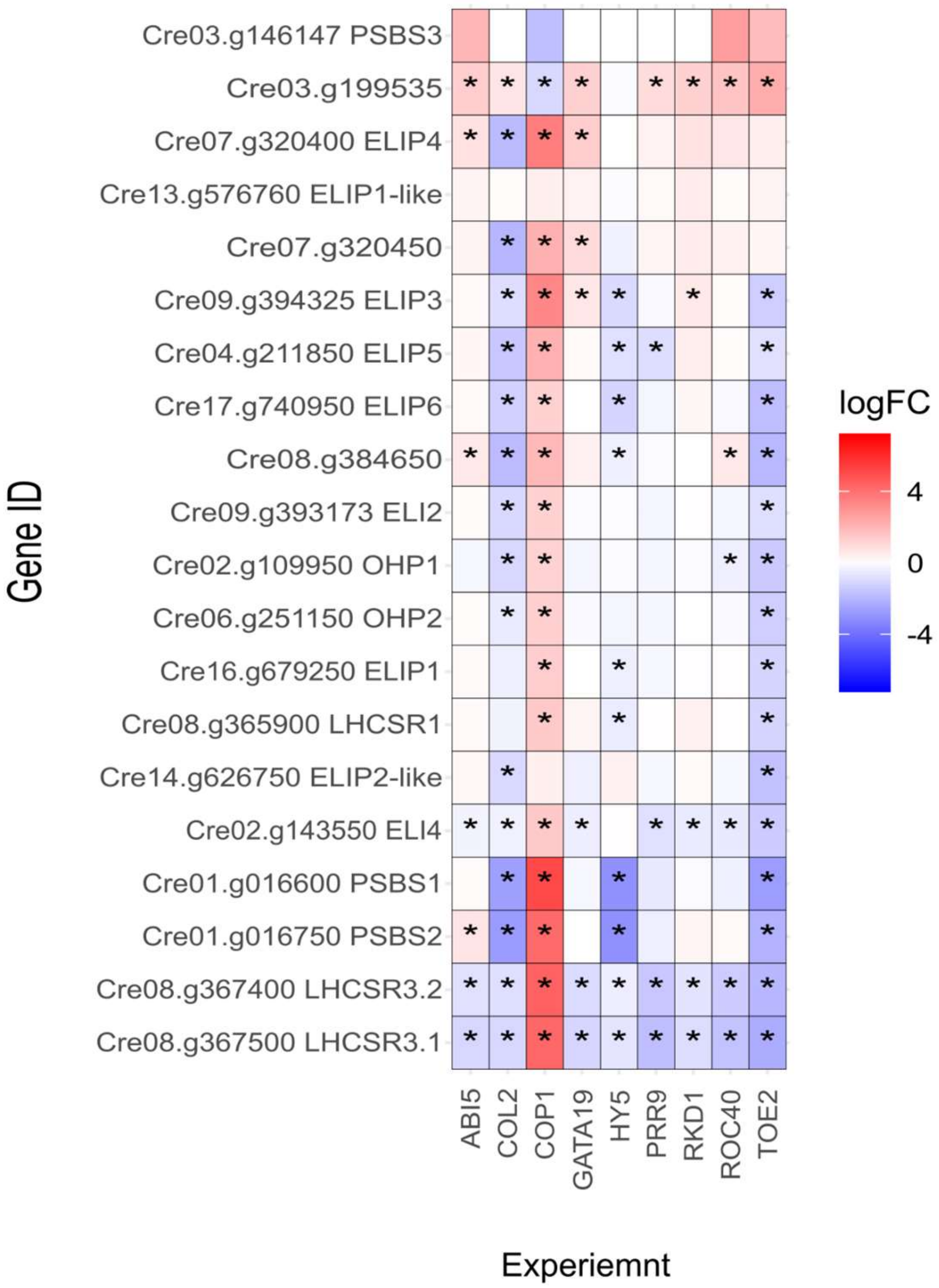


*Figure S5: Heatmap (red upregulated, blue downregulated) of the photoprotection genes (PP) for the mutants abi5, col2, cop1 gata19, hy5, prr9, rkd1, roc40 and toe2. Significantly differential genes p<0.01 are marked (*).*

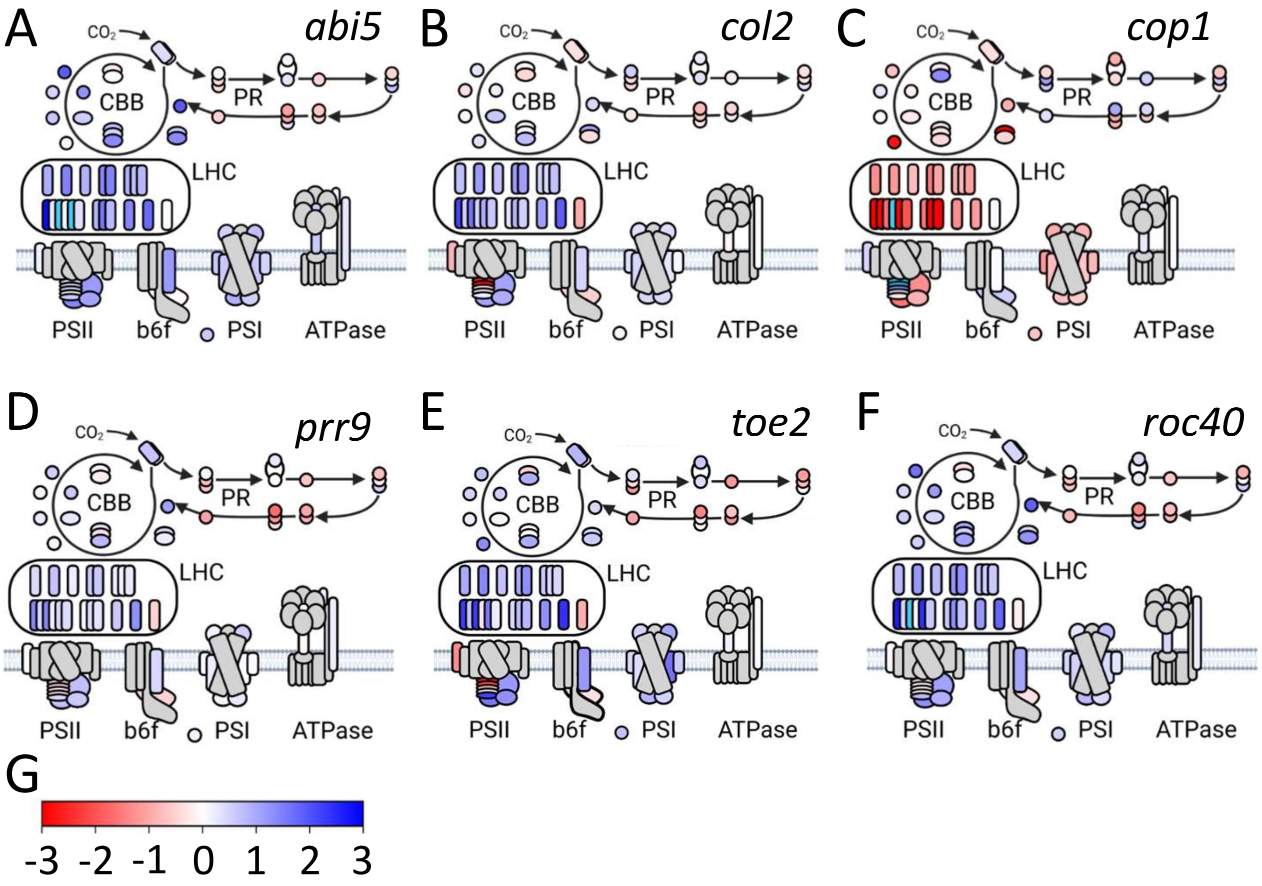


*Figure S6: A-I: Photosynthesis pathway heatmap of* abi5 *(A),* col2 *(B),* cop1 *(C), prr9 (D),* toe2 *(E) and* roc40 *of the logFC (-3 red, 0 white, 3 blue) of the CBB, PR, LHC, PSII, b6f, PSI and ATPase. Nuclear encoded genes are grey and genes with a logFC>3 are in light blue and with a logFC<-3 in pink. G: color scale for the heatmaps A-I of the logFC (-3 red, 0 white, 3 blue)*

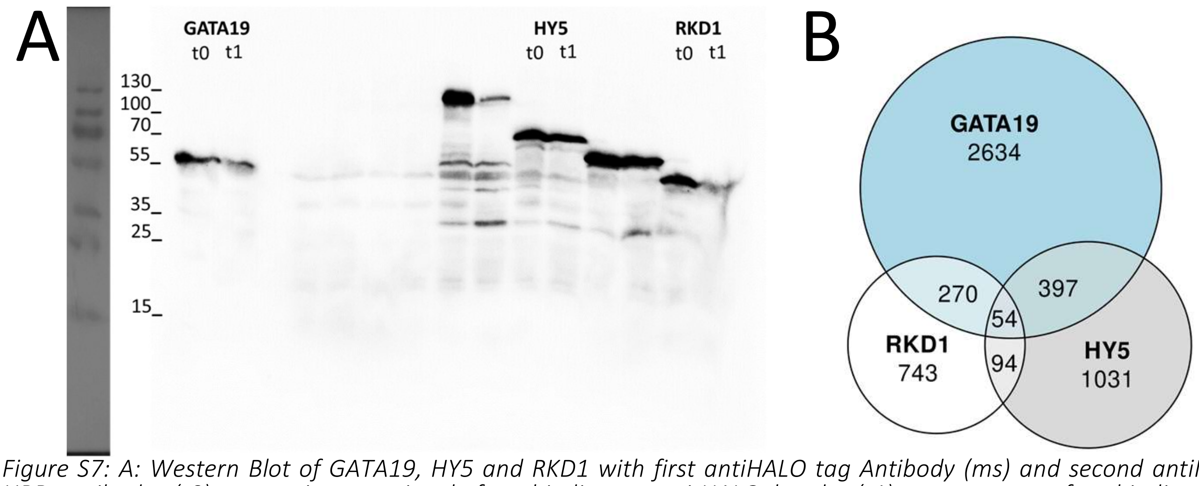


*Figure S7: A: Western Blot of GATA19, HY5 and RKD1 with first antiHALO tag Antibody (ms) and second antiMS HRP antibody. (t0) expression reaction before binding to anitHALO beads, (t1) supernatant after binding to antiHALO beads. B: Venn diagram of bound genes in ampDAPseq of GATA19, RKD1 and HY5. Areas are proportional to the size of the group. Number of peaks detected were 23854 for GATA19, 20156 for HY5 and 6142 for RKD1.*

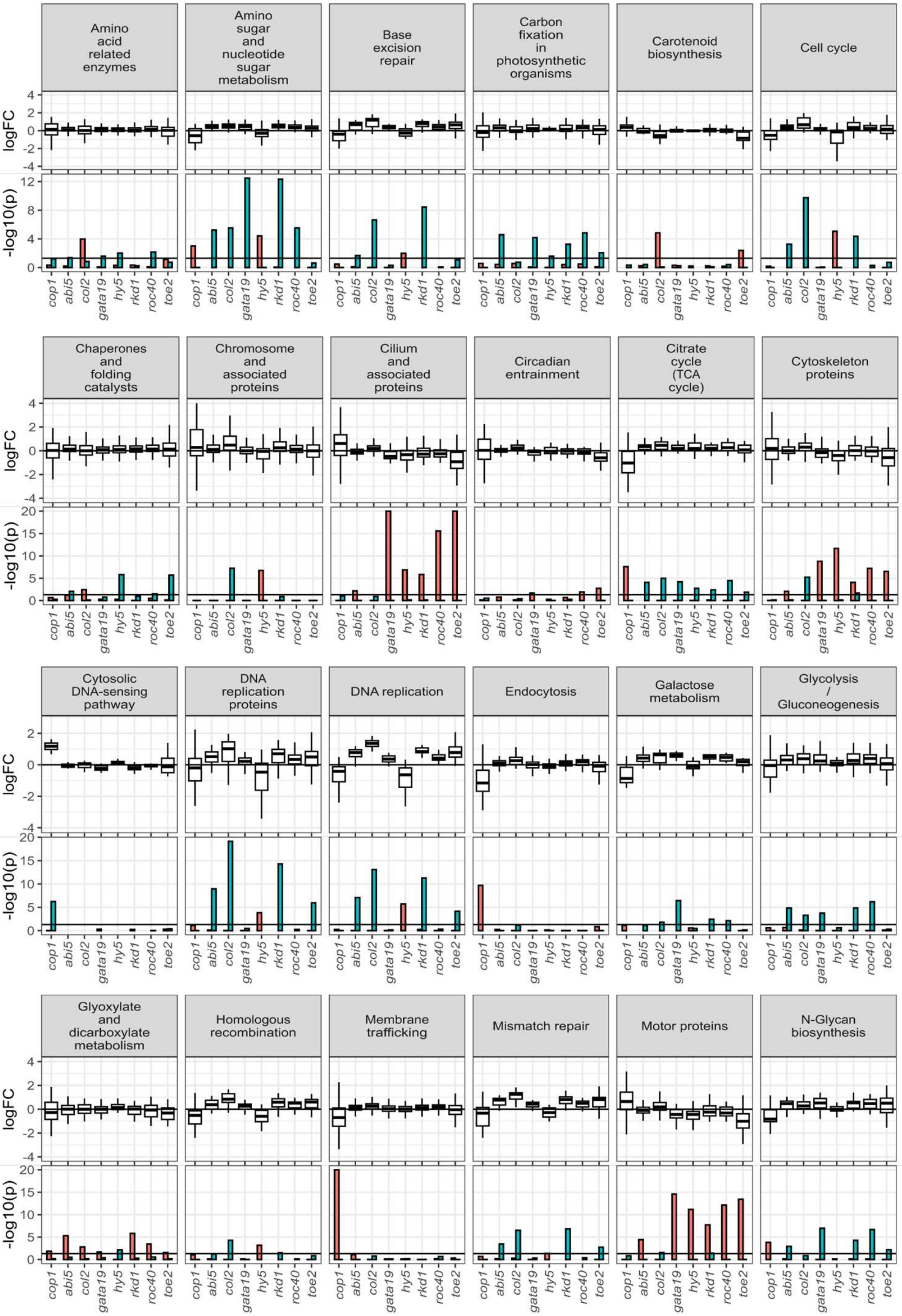

Amino acid related enzymes
Amino sugar and nucleotide sugar metabolism
Base excision repair
Carbon fixation in photosynthetic organisms
Carotenoid biosynthesis
Cell cycle
Chaperones and folding catalysts
Chromosome and associated proteins
Cilium and associated proteins
Circadian entrainment
Citrate cycle (TCA cycle)
Cytoskeleton proteins
Cytosolic DNA-sensing pathway
DNA replication proteins
DNA replication
Endocytosis
Galactose metabolism
Glycolysis / Gluconeogenesis
Glyoxylate and dicarboxylate metabolism
Homologous recombination
Membrane trafficking
Mismatch repair
Motor proteins
N-Glycan biosynthesis
logFC
-log10(p)
cop1
abi5
col2
gata19
hy5
rkd1
roc40
toe2

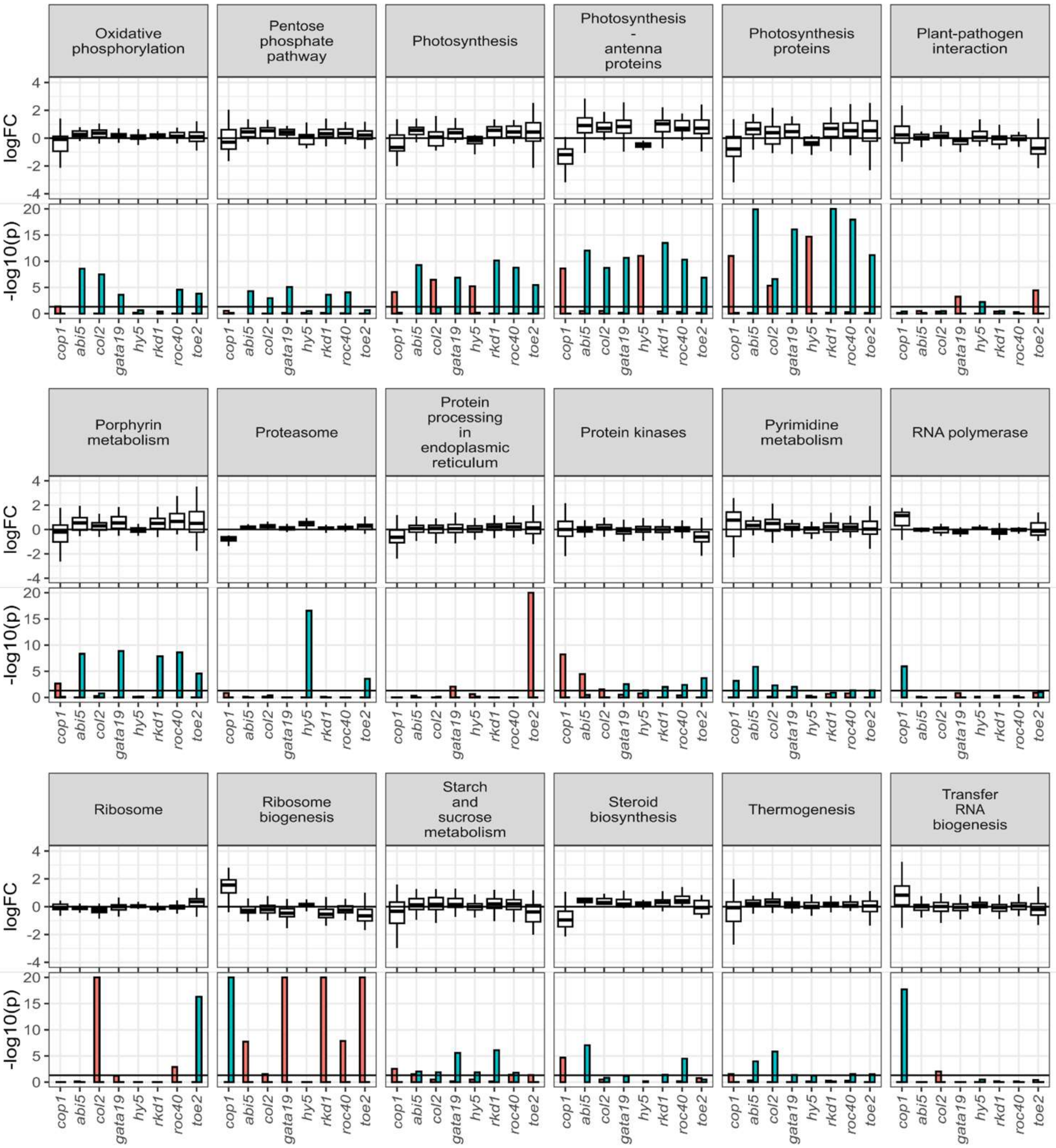


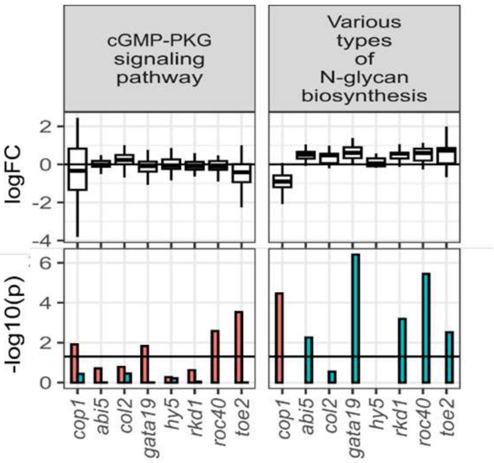


*Figure S8: Top panel boxplot of the logFC and jitter of individual genes of the mutants cop1, abi5, col2, gata19, hy5, prr9, rkd1, roc40 and toe2. The bottom panel bar plot of the –log10 p-value of the gene list enrichment of the significantly upregulated genes (green) and significantly downregulated genes (red).*

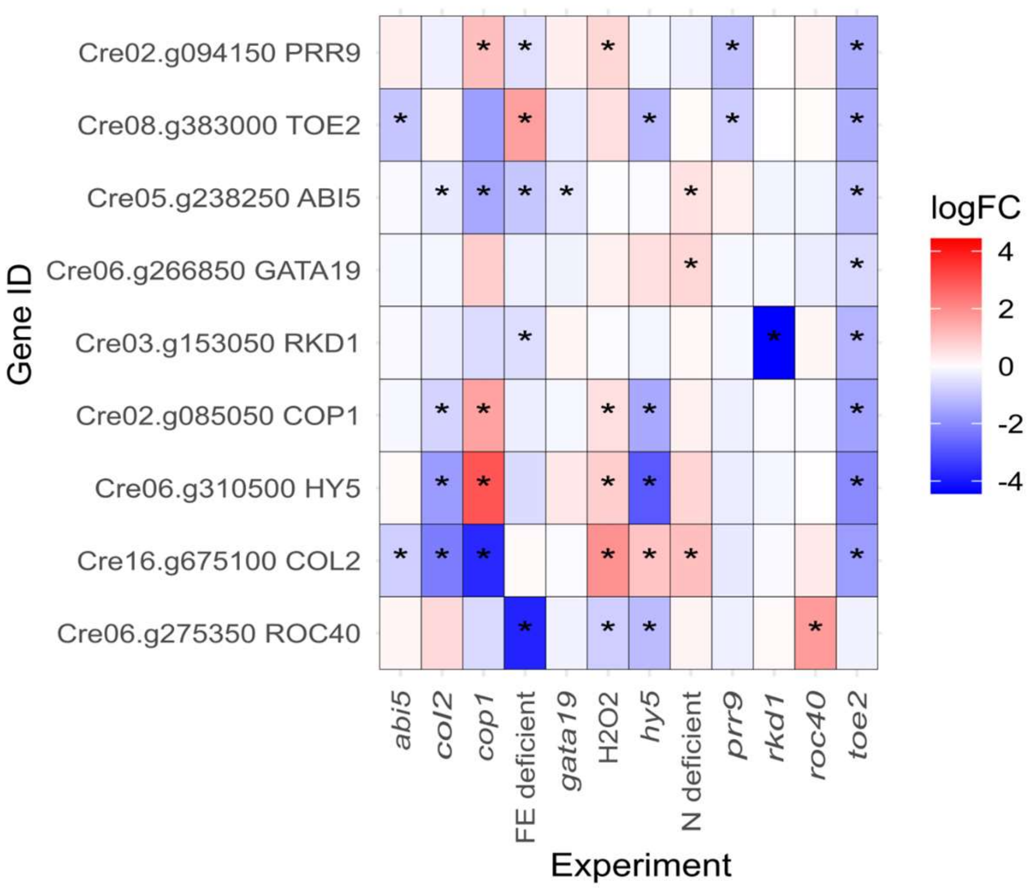


*Figure S9: Heatmap (red upregulated, blue downregulated) of the transcription factors ABI5, COL2, COP1 GATA19, HY5, PRR9, RKD1, ROC40 and TOE2 for the mutants abi5, col2, cop1 gata19, hy5, prr9, rkd1, roc40, toe2, H2O2 under iron and nitrogen deficiency. Significantly differential genes p<0.01 are marked (*).*
